# Acute *FMR1* reduction alters cellular and circuit properties in human cortex

**DOI:** 10.64898/2026.03.11.711123

**Authors:** Aditi Singh, Saman Abbaspoor, Leeyup Chung, Maxwell J. Heinrich, Scellig Stone, Hart Lidov, Beatriz Maio, Tien Phuoc Tran, Jaeyoung Yoon, Jiawen Teng, Catalina Martinez-Reyes, Elijah Hammarlund, Xiguang Xu, Remi Hirsh, Ryan Guardado, Elizabeth Buttermore, Alexander Rotenberg, Jeff Gavornik, Brielle Ferguson, Jordan S. Farrell, Emily K. Osterweil

## Abstract

Transcriptional silencing of *FMR1* results in Fragile X syndrome (FXS), the leading inherited cause of intellectual disability (ID) and autism. Although the impact of *FMR1* reduction on cortical function has been studied using the *Fmr1^-/y^* mouse model, mechanistic insights from the human brain are lacking. Here we show that acute *FMR1* reduction in *ex vivo* human cortical slices induces transcriptomic changes that are concordant with those observed in FXS patient cortex, identifying many targets not altered in the *Fmr1^-/y^* mouse model. Consistent with changes in ion channel expression specifically in human tissue, *FMR1* reduction drives increased intrinsic excitability of deep-layer pyramidal neurons and network-level population bursts observed with two-photon calcium imaging. Together, these findings define the consequences of *FMR1* loss in human cortex and establish an experimental framework for identifying human-specific mechanisms and therapeutic targets relevant to FXS.

## Main Text

Fragile X Syndrome (FXS) is a neurodevelopmental disorder caused by transcriptional silencing of the *FMR1* gene and subsequent loss of the RNA binding protein Fragile X Messenger Ribonucleoprotein (FMRP). In the brain, FMRP represses mRNA translation ^1–3^, regulates RNA stability^4,5^, and is involved with trafficking RNA to synapses ^6^. FMRP has also been implicated in ion channel regulation through direct binding ^7–9^. Mechanistic exploration of the *Fmr1* KO (*Fmr1^-/y^*) mouse cortex reveals a complex picture of altered cellular and network level brain function following FMRP loss, including hyperexcitability in some circuits ^10,11^, aberrant excitatory and inhibitory synaptic connectivity ^12–16^, and augmented synaptic plasticity ^17–20^. In contrast, how *FMR1* loss affects human cortex is not known.

With no targeted therapies to alleviate major symptoms in FXS patients, understanding the relevance of FMRP to human brain circuits is crucial, especially if *FMR1* reduction induces changes in the human brain that are absent or muted in mouse. Indeed, recent studies highlight interspecies differences in cortical neuronal structure and connectivity ^21,22^, synaptic transmission ^22–24^, and cell type diversity^25,26^, which may impinge on unique effects of *FMR1* reduction in humans. Together with evidence that FMRP associates with additional mRNA transcripts in human neurons ^27^, these species differences suggest that *FMR1* reduction may produce distinct molecular and functional consequences in human brain. We therefore investigated the outcome of acutely reducing *FMR1* expression in human cortex.

### Reduction of *FMR1* in human cortex

To determine how *FMR1* reduction alters the human cortex across molecular and functional assays, we first established a platform to reduce *FMR1* expression in human cortical slices. To preserve the full suite of cortical cell types, their integration into cortical circuits, and tissue health across weeks of investigation, we employed an organotypic slice culture strategy prepared from non-lesional cortex collected during pediatric epilepsy surgery (see Methods) ^28–30^ and applied AAVs to reduce *FMR1* expression using shRNA (**Fig. 1a, Extended Data Fig. 1a-b and Supplementary Table 1**). Crucially, we utilized a within-resection design to treat interleaved slices with *FMR1* shRNA (sh*FMR1*) or scrambled shRNA (shScr) for every resection to overcome the challenge of patient-to-patient variability. By 7 days in culture, *FMR1* mRNA is reduced by >80% in FACS-sorted transduced sh*FMR1* neurons as compared to shScr neurons (**Fig. 1b**) and FMRP protein levels per neuron are reduced by >50% (**Fig. 1c and Extended Data Fig. 1c-d**). While the acute nature of this approach is unlikely to capture neurodevelopmental changes, previous work shows acute *FMR1* reduction in adult neurons recapitulates multiple molecular and circuit-level FXS phenotypes across models^27,31–33^, suggesting that many symptoms are due to the ongoing loss of FMRP. Supporting this, FMRP replacement studies in mice show that *FMR1* reintroduction resolves phenotypes even when administered in adult brain^34–36^.

**Figure 1.**
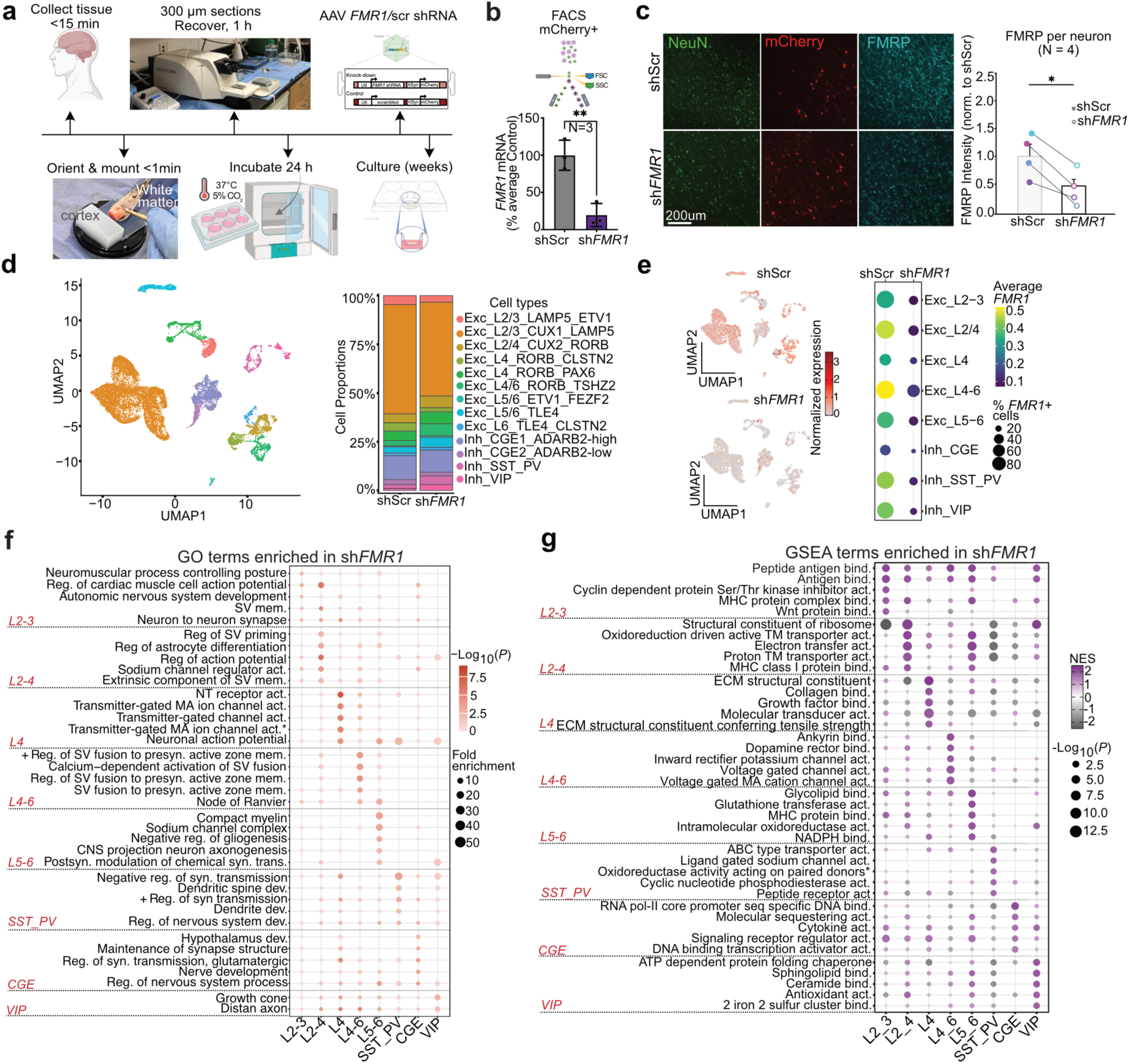
Reduction of *FMR1* in human cortex causes distinct molecular changes in cortical neurons. **(a)** Schematic illustrates the experimental timeline for cortical slice preparation, transduction and analysis. **(b)** FACS of mCherry+ neurons was performed on shScr and sh*FMR1* expressing slices at 7 days post-transduction. *FMR1* mRNA levels in samples transduced with sh*FMR1* or scrambled control (shScr). Points represent independent patients (N = 3); bars show the group mean with error bars, (Mean difference = −80.31 ± 6.95, paired t-test, *P* = 0.0025). **(c)** Representative ROI of shScr and sh*FMR1* staining for NeuN, mCherry, and FMRP. scale=200um. Quantification revealed a significant reduction in FMRP intensity per neuron (normalized to shScr; N = 4 patients) in shRNA-treated cortical slices compared to shScr controls. Each color represents neurons from the same slice/patient, (Mean difference = −63.5 ± 0.31, paired t-test, *P* = 0.0268). **(d)** UMAP visualization and cell-type composition (right) across excitatory and inhibitory classes show cellular subtypes present in both shScr and sh*FMR1*-expressing neurons. **e)** Distribution and expression levels of *FMR1* across treatment conditions. Feature plots (left) show normalized *FMR1* expression density mapped onto UMAP space and dot plot quantifies the average expression level (color) and percentage of cells expressing *FMR1* (dot size) across resolved excitatory and inhibitory cell types, confirming robust knockdown. **(f)** Dot plot shows the top 5 nervous-system-related GO terms (*P* < 0.01, ≥ 3 genes, ranked by fold enrichment) enriched among differentially expressed genes (pseudobulk DESeq2, sh*FMR1* vs shScr, *P* < 0.01) in each cell type; each term is shown for all cell types (dot size = Fold enrichment; dot color = −Log10(*P*)). *Transmitter-gated monoatomic ion channel activity involved in regulation of postsynaptic membrane potential. **(g)** Dot plot shows the top 5 GSEA molecular function terms enriched in sh*FMR1* neurons (*P* < 0.01, highest positive NES) for each cell type (genes ranked by the DESeq2 Wald statistic); dot color indicates normalized enrichment score (NES) and dot size indicates −Log10(*P*). *Oxidoreductase activity acting on paired donors, with incorporation or reduction of molecular oxygen, NAD(P)H as one donor, and incorporation of one atom of oxygen. Abbreviations (f, g): act., activity; bind., binding; dev., development; ECM, extracellular matrix; MA, monoatomic; mem., membrane; NT, neurotransmitter; pol-II, polymerase II; postsyn., postsynaptic; presyn., presynaptic; Reg., regulation; +Reg., positive regulation; seq, sequence; SV, synaptic vesicle; syn., synaptic; TM, transmembrane; trans., transmission. Panel b-c: \**P* < 0.05, \*\**P* < 0.01.

To assess the transcriptomic impact of *FMR1* reduction in human cortex, we performed single cell (sc)RNA-sequencing on multiple pairs of sh*FMR1* and shScr slices. We utilized a whole-cell approach rather than single-nucleus RNA sequencing (snRNA-seq) to ensure comprehensive recovery of the cytoplasmic transcriptome, where FMRP functions as an RNA-binding protein to regulate transcript stability and translation. We used fluorescence activated cell sorting (FACS) to isolated neurons based on mCherry fluorescence driven by a synapsin promoter and prepared single-cell libraries using 10X genomics Chromium GEM-X, which obtained >75% viability with only DAPI-/DRAQ+ neurons included in our analyses (**Extended Data Fig. 1e-f**). Multiple neuronal clusters were annotated using reference-based mapping to an age-matched human mid-temporal gyrus atlas^37^ and canonical cell-type marker expression (**Extended Data Fig. 2a**). Identified clusters include Excitatory neurons from L2/3, L2/4, L4, and L5/6, along with GABAergic parvalbumin (PV), somatostatin (SST), vasoactive peptide (VIP) subclasses, as well as a larger immature Caudal Ganglionic Eminence (CGE)-derived subclass that may be present due to the young age of donating patients (10 years old). Overall cell-type representation was comparable across shScr and sh*FMR1* conditions (**Fig. 1d and Extended Data Fig. 2b**), enabling comparisons of gene expression within neuronal subtypes. *FMR1* mean expression level and percentage of *FMR1*-containing cells is substantially reduced in every sh*FMR1* subtype examined (**Fig. 1e**). In addition, FMRP target transcripts are consistently downregulated in sh*FMR1* neurons (**Extended Data Fig. 3a, Supplementary Table 2**), consistent with FMRP loss ^38–40^.

### *FMR1* reduction alters gene expression across cortical layers

To identify transcriptomic changes induced with *FMR1* reduction, we performed cell-type resolved pseudobulk differential expression analyses between sh*FMR1* and shScr populations (**Extended Data Fig. 3b, Supplementary Table 3**). To determine the pathways altered within each neuronal subtype, we performed Gene Ontology (GO) analysis on the most significantly changed transcripts, and Gene Set Enrichment Analyses (GSEA) of the full ranked gene lists to capture smaller, coordinated shifts in functional categories (**Fig. 1f-g, Supplementary Table 4-5**). GO analyses show an enrichment of terms related to synaptic function across cell types. In upper L2-3 and L2-4 neurons, changes in presynaptic vesicle cycle and peptide/antigen binding are observed, while those related to neurotransmitter receptor function are observed in L4. In deeper L4-6 and L5-6 neurons, changes in synaptic vesicle release are accompanied by altered ion channel function as well as glial terms. Synaptic terms also dominate the changes in all interneuron subtypes. GSEA highlights additional changes in pathways related to ribosomal function and oxidative phosphorylation, consistent with a loss in FMRP, which are up- or downregulated in a subtype-specific manner.,. This analysis also highlights changes related to ion channel, which are concentrated in L4-6 neurons (13 terms). Together, these results suggest loss of *FMR1* results in a dysregulation of synaptic and metabolic function across all neuronal subtypes.

### FXS patient cortex exhibits greater transcriptomic overlap with *FMR1*-deficient human slices than *Fmr1^-/y^* mouse cortex

To investigate the degree to which transcriptomic changes in *FMR1*-deficient human slices recapitulate changes in FXS patient cortex, we performed a series of overlap analyses comparing our results to snRNA-seq data from FXS patient versus control cortex^41^ (**Fig. 2a-e, Supplementary Table 7-8**). To benchmark this comparison, we performed the same comparative analyses between *FMR1*-deficient human slices and scRNA-seq data from *Fmr1^-/y^*versus wild-type mouse cortex^38^. Notably, like FXS patients, *Fmr1^-/y^* mice exhibit *FMR1* loss *in utero*. To compare across groups with different sample sizes and cell type representations, we collapsed cell types into broad excitatory and inhibitory subgroups (**Supplementary Table 6)**.

**Figure 2.**
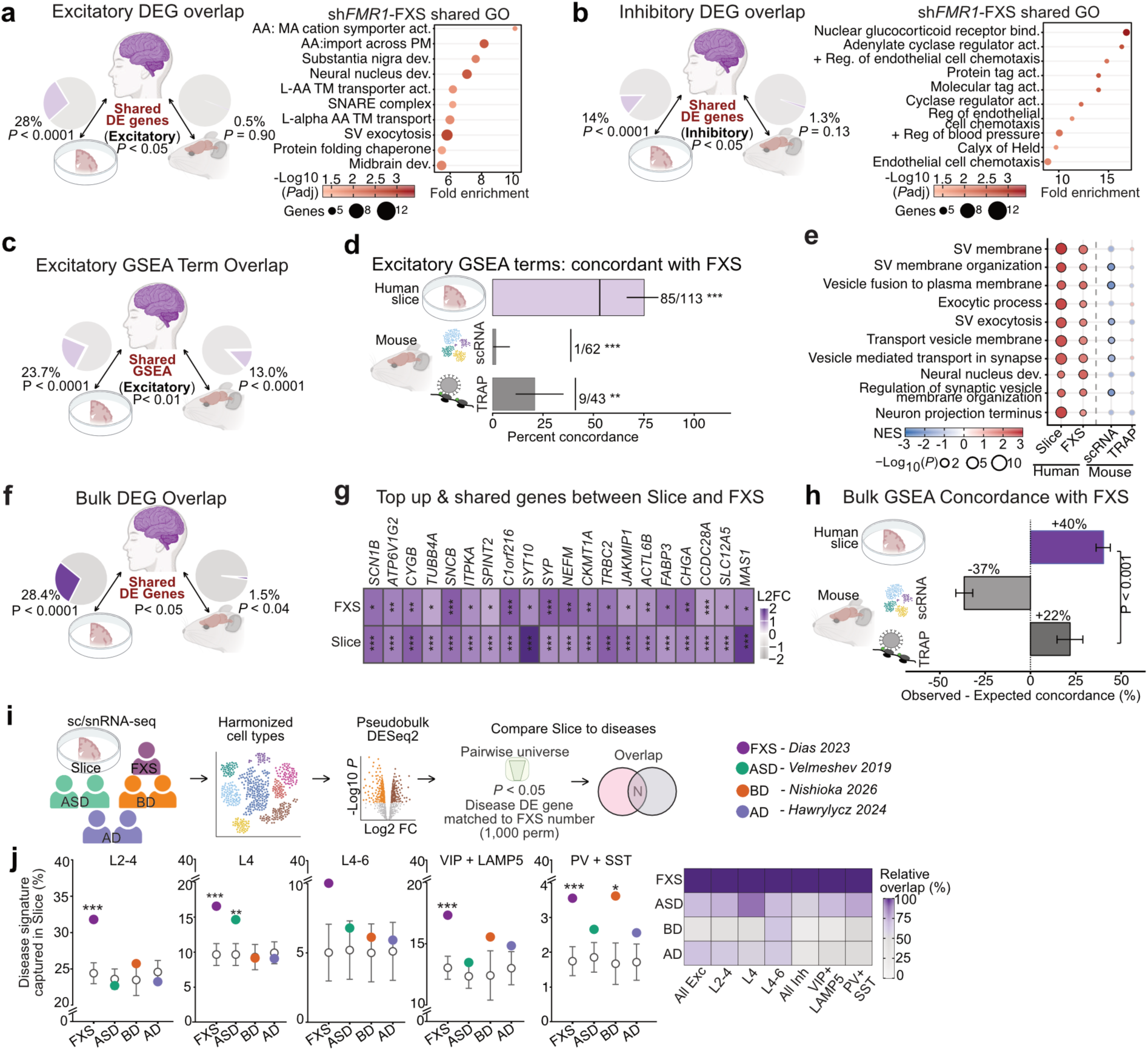
Shared cell-type transcriptomic signatures in *FMR1* deficient human cortex and FXS. (**a-b**) Overlap of differentially expressed genes (DEGs; nominal *P* < 0.05) of FXS patients (snRNA-seq) with sh*FMR1* human slices (scRNA-seq; left) and P5 *Fmr1*-KO mouse (scRNA-seq; right) in excitatory **(a)** and inhibitory **(b)** neurons. Pies show the percentage of FXS DEGs shared, among genes tested in both datasets; *P*, permutation test. Dot plots show the top 10 GO terms, ranked by fold enrichment, among genes with *P* < 0.05 in both FXS patients and sh*FMR1* slices (adjusted *P* < 0.05, ≥ 5 genes). Dot color indicates −Log10(adjusted *P*); dot size, number of genes. **(c)** Overlap of GSEA GO terms (*P* < 0.01) of FXS patients with human slices (left) and P5 mouse (right) in excitatory neurons; percentages of FXS terms; P, one-sided Fisher’s exact test. **(d)** Percentage of GSEA terms significant (*P* < 0.01) in both FXS patients and each dataset (human slice; P5 mouse scRNA-seq; adult mouse Camk2a-TRAP-seq) with the same NES direction, in excitatory neurons. Labels, concordant/jointly significant terms; error bars, 95% Wilson confidence interval; vertical line, concordance expected by chance; two-sided binomial test. **(e)** Top 10 GSEA terms significant (*P* < 0.01) and concordant in human slices and FXS patients, ranked by mean NES, shown for all four datasets. Dot color, normalized enrichment score (NES); dot size, - Log10(P); outlined, *P* < 0.01. **(f)** Overlap of DEGs (*P* < 0.05) of FXS patient cortex (bulk RNA-seq; GSE296077, 5 FXS vs 5 typically developing donors) with sh*FMR1* human slices (left) and P5 *Fmr1*-KO mouse (right), all neurons combined (excitatory and inhibitory neurons pooled into one pseudobulk per donor or animal). Pies show the percentage of FXS bulk DEGs shared, among genes tested in both datasets; P, permutation test. **(g)** Top 20 genes with *P* < 0.05 in both FXS bulk cortex and sh*FMR1* slices and increased in both, ranked by slice P (left to right). Cell color, log2 fold change; \**P* < 0.05, \*\**P* < 0.01, \*\*\**P* < 0.001. **(h)** Direction concordance of GSEA GO terms with FXS bulk cortex, corrected for chance, for human slices, P5 mouse scRNA-seq and adult mouse TRAP-seq (Camk2a- and Pvalb-TRAP combined), all neurons combined. Bars, percentage of terms significant (*P* < 0.01) in both datasets with the same NES direction, minus the percentage expected by chance (from each dataset’s proportion of positive-NES terms). Error bars, 95% confidence interval and bracket, two-sided P for the slice versus TRAP difference, both from 5,000 bootstrap resamples of GO terms. **(i)** Schematic of the disease-overlap mapping pipeline: harmonization of single-cell/single-nucleus datasets into corresponding cell-type groups, pseudobulk differential expression (DE) analysis and pairwise comparison of sh*FMR1* slice DE genes with disease DE genes. **(j)** Percentage of disease DEGs (*P* < 0.05) also DE in sh*FMR1* slices (*P* < 0.05), among genes tested in both, for Fragile X Syndrome (FXS; Dias 2023), Autism Spectrum Disorder (ASD; Velmeshev 2019), Bipolar Disorder (BD; Nishioka 2026) and Alzheimer’s Disease (AD; Hawrylycz 2024). ASD, BD and AD DEGs were down-sampled to the number of FXS DEGs (mean of 1,000 draws); open circles, mean ± s.d. of 10,000 size-matched random gene sets; broken y axes where indicated. P versus background, corrected across 28 tests with the Benjamini–Hochberg method; *Padj < 0.05, **Padj < 0.01, ***Padj < 0.001. Heatmap, overlap relative to FXS (FXS = 100%). Abbreviations **(a, b, e)**: AA, amino acid; act., activity; bind., binding; dev., development; MA, monoatomic; PM, plasma membrane; Reg., regulation; +Reg., positive regulation; SV, synaptic vesicle; TM, transmembrane.

Despite the differences in patient age and sample preparation, we find that 28% of the DEGs identified in FXS patient excitatory neurons are also significantly altered in *FMR1*-deficient excitatory neurons from human slices, which is significantly greater than expected (**Fig. 2a, Extended Data Fig. 4a, c**). Interestingly, the equivalent overlap between *Fmr1^-/y^* mouse cortex and FXS patient cortex is smaller, with only 0.5% DEGs shared between excitatory neuron populations. For inhibitory neurons, a significant 14% overlap in DEGs is observed between *FMR1*-deficient slices and FXS patients, which is also larger than the 1.3% overlap seen between FXS patients and the *Fmr1^-/y^*cortex (**Fig. 2b, Extended Data Fig. 4b, d**). This shows the transcriptomic profile of *FMR1*-deficient neurons overlaps with that of FXS cortex to a greater extent than of the *Fmr1^-/y^* cortex (**Supplementary Table 7, Extended Data Fig. 4a-d**). Genes altered in both FXS patients and *FMR1*-deficient slices were enriched for GO terms related to synaptic vesicle exocytosis and the SNARE complex in excitatory neurons (**Fig. 2a, b; Supplementary Table 9**).

The difference in DEG overlap between sh*FMR1* human slices and the *Fmr1^-/y^* mouse led us to investigate whether the same is seen in GSEA categories, which are less reliant on individual gene changes (**Fig. 2c and Extended Data Fig. 4e-f, Supplementary Table 10**). We find FXS patient and *FMR1*-deficient human slices share 23.7% of differentially expressed terms in excitatory neurons, which is greater than the 13.0% seen in *Fmr1^-/y^*mice. Additionally, we performed a concordance analysis to determine whether the overlapping categories are changed in a similar direction (**Fig. 2d; Extended Data Fig. 4g–l**). 75.2% of GSEA terms identified in FXS patient cortex are concordant in the sh*FMR1* human slice, whereas only 1.6% are concordant with the *Fmr1^-/y^* mouse. To explore this result further, we examined the concordance of GSEA terms seen in the translating ribosome (TRAP) fraction of *Fmr1^-/y^* Camk2a excitatory and PV inhibitory neurons recently published in Suresh et al.^42^. Here there was greater concordance with FXS patients (20.9%) than observed with scRNA-seq, but still much lower than the 75.2% seen with sh*FMR1* human slices. Thus, more differentially altered pathways are shared between sh*FMR1* human slices and FXS patients, and of those shared pathways, more are altered in a similar direction (**Fig. 2e**). Since these results were largely mirrored in inhibitory cell types (**Extended Data Fig. 4e–l**), together these data suggest that changes in sh*FMR1* human may be a more accurate reflection of transcriptomic changes in FXS than the *Fmr1^-/y^* mouse.

To test whether this correspondence extends to an independent FXS patient cohort, we repeated these analyses against a recently published bulk RNA-sequencing dataset from post-mortem cortex of five FXS and five age-matched typically developing donors^43^, which shares no donors with the snRNA-seq cohort analyzed above. Because this dataset profiles whole cortical tissue rather than sorted cell types, we pooled excitatory and inhibitory neurons into a single pseudobulk profile per donor or animal for each comparator. Of the genes differentially expressed in this second FXS patient cohort, 28.4% are also differentially expressed in *FMR1*-deficient human slices (*P* < 0.0001), compared with 1.5% in the *Fmr1^-/y^*mouse (*P* = 0.04) (**Fig. 2f**). The genes altered in both the FXS patient cortex and *FMR1*-deficient slices include *SCN1B* together with further transcripts encoding ion channel, synaptic vesicle and cytoskeletal proteins (**Fig. 2g**). At the pathway level, GSEA terms significant in both datasets are concordant in direction 40 percentage points above chance in *FMR1*-deficient human slices, whereas the P5 *Fmr1^-/y^* scRNA-seq dataset lies 37 percentage points below chance and adult *Fmr1^-/y^* TRAP-seq 22 percentage points above it (*P* < 0.001; **Fig. 2h**). The closer correspondence between *FMR1*-deficient human slices and FXS patient cortex is therefore reproduced in a second patient cohort assayed by a different method.

### Reduction of *FMR1* recapitulates FXS-specific changes

One potential explanation for the greater overlap and concordance of sh*FMR1* human slices and FXS patients is that the data are obtained from the same species. Thus, rather than specifically reflecting loss of FMRP, the shared DEG and pathway changes could reflect general alterations in neuronal function that would be expected to occur across other neurological and psychiatric disorders. To that end, we compared the overlap of cell-type-specific transcriptomic changes in *FMR1*-deficient slices to those in multiple human brain disorders including Autism Spectrum Disorder (ASD)^44^, Bipolar Disorder (BD)^45^, and Alzheimer’s Disease (AD)^46^, comparing to the overlap seen with FXS^41^ (**Fig. 2i-j**). To account for differences in sample size between single-cell datasets, we calculated DEGs from harmonized cell types using permutations of the same number of groups for each disorder, then compared the overlapping proportion of DEG in sh*FMR1* slices to each mean (**Extended Data Fig. 5a-b**, Methods). The degree of overlap between changes in sh*FMR1* slices and FXS is significantly greater than chance in almost all cell types. In contrast, no significant overlap is seen with AD, and only one subtype shows a significant overlap with ASD (L4) or BD (PV + SST) (**Fig. 2i-j**). Together, these results support that the similarity of changes between FXS and sh*FMR1* slices is not explained by species conservation and are therefore more reflective of FXS neurobiology.

### *FMR1* reduction increases intrinsic excitability in human cortical pyramidal neurons

Among the shared changes we identified in excitatory *FMR1*-deficient cortex and FXS patients are genes encoding ion channel subunits involved in maintaining neuronal excitability (**Extended Data Fig. 3b; Extended Data Fig. 4c, d; Supplementary Table 7**). Although previous work has demonstrated region and layer-specific changes in intrinsic excitability in *Fmr1^-/y^*mouse cortex, the effect of *FMR1* reduction in human cortex remains unknown. To establish this, we performed whole cell patch-clamp recordings on pyramidal neurons in sh*FMR1*- and shScr-treated slice pairs taken from 10 patients (**Supplementary Table 1**). Hyperpolarizing and depolarizing step currents were injected into current-clamped mCherry-expressing deep layer pyramidal neurons with biocytin included in the pipette for *post hoc* confirmation of laminar depth and morphology (**Fig. 3a**).

**Figure 3.**
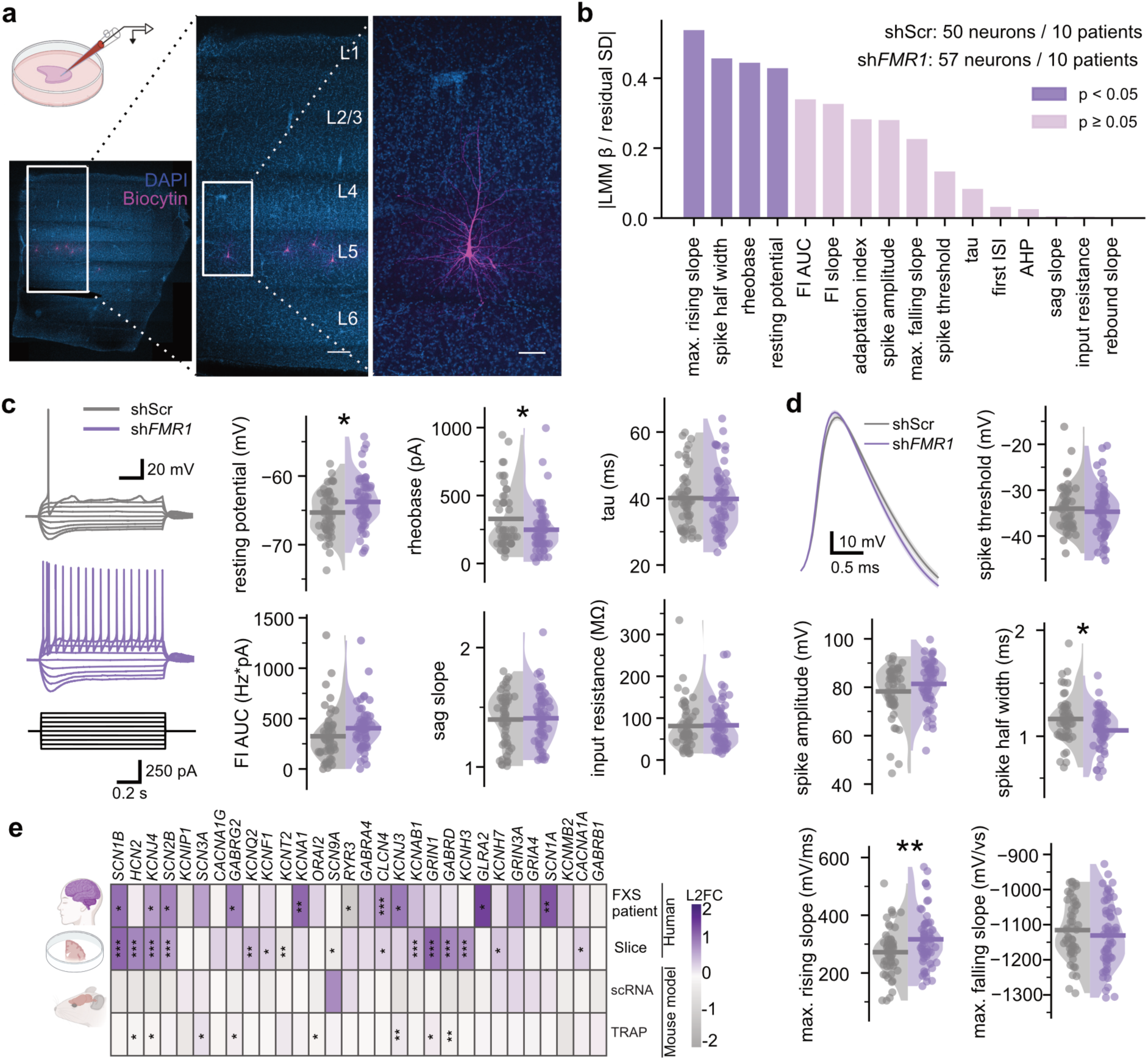
*FMR1* reduction elevates intrinsic excitability in human cortical pyramidal neurons. **(a)** Schematic of whole-cell electrophysiology performed on pyramidal neurons (top left). Biocytin (magenta) included in the whole-cell internal solution of each recorded neuron was visualized, revealing laminar position and morphology**. (b)** Rank-ordered linear mixed-effects model (GLMM, see *Methods & Materials*) effect sizes for each quantified electrophysiological metric. **(c)** 1-second hyperpolarizing and depolarizing step currents (−200 to 1000 pA in 50 pA increments) were injected into neurons from scrambled- (shScr, top) and *FMR1*-knockdown-treated (sh*FMR1*, bottom) neurons, where sh*FMR1*-treated neurons exhibit depolarized resting potential (shScr: −65.3 ± 0.5 mV (n=50), sh*FMR1*: −63.8 ± 0.5 mV (n=57), *P* = 0.03), reduced rheobase current (shScr: 328 ± 32 pA (n=50), sh*FMR1*: 248 ± 23 pA (n=57), *P* = 0.02) and normal adaptation index (shScr: 1.59 ± 0.18, sh*FMR1*: 1.29 ± 0.09 , *P* = 0.15), sag slope (shScr: 1.40 ± 0.03 (n=50), sh*FMR1*: 1.41 ± 0.03 (n=54), *P* = 0.98), FI slope (shScr: 0.062 ± 0.006 Hz/pA (n=43), sh*FMR1*: 0.073 ± 0.006 Hz/pA (n=52), *P* = 0.12) and FI area under the curve (scr: 325 ± 36 Hz·pA (n=50), sh*FMR1*: 404 ± 29 Hz·pA (n=57), *P* = 0.08). **(d)** Active properties were extracted from the first action potential generated by the step currents in (c), revealing reduced spike half-width (shScr: 1.16 ± 0.03 ms (n=50), sh*FMR1*: 1.07 ± 0.03 ms (n=57), *P* = 0.02), increased maximum rising slope (shScr: 272 ± 12 mV/ms (n=50), sh*FMR1*: 317 ± 13 mV/ms (n=57), *P* = 0.006), and normal maximum falling slope (shScr: −1115 ± 12 mV/ms (n=50), sh*FMR1*: −1131 ± 12 mV/ms (n=57), *P* = 0.25), spike threshold (shScr: −34.0 ± 0.7 mV (n=50), sh*FMR1*: −34.7 ± 0.7 mV (n=57), *P* = 0.50) and spike amplitude (shScr: 78.3 ± 1.5 mV (n=50), sh*FMR1*: 81.4 ± 1.3 mV (n=57), *P* = 0.15). * *P* < 0.05, ** *P* < 0.01, linear mixed-effects model with shRNA as main factor. **(e)** Heatmap of log2 fold change (L2FC) of ion-channel genes in excitatory neurons of FXS patients (FXS vs control), sh*FMR1* slices (sh*FMR1* vs shScr), P5 *Fmr1*-KO mouse scRNA-seq and adult *Fmr1*-KO mouse Camk2a-TRAP-seq (KO vs WT; mouse genes mapped to human 1:1 orthologs). DESeq2 Wald test (nominal *P*), \**P* < 0.05, \*\**P* < 0.01., \*\*\**P* < 0.001.

To control for patient age, resection location, and other variables, all data were collected using the same paired design (sh*FMR1* or shScr treatment in adjacent slices) and analyzed using a linear mixed model (LMM) (**Supplementary Table 11**). Notably, analysis of 107 deep layer pyramidal neurons across 10 patients revealed increased excitability of frontal cortex neurons compared to those from temporal cortex and several age-dependent properties, including increased *h*-current-mediated sag and rebound slopes^47^ (**Extended Data Figs. 7-8, Supplementary Table 11**). Despite the variance introduced by these factors, *FMR1* had a significant impact on several intrinsic properties consistent with increased excitability. Namely, sh*FMR1*-treated neurons exhibit significantly depolarized resting potentials and require less current to generate an action potential compared to those treated with shScr (**Fig. 3b-c, Extended Data Fig. 6**). We further investigated active and passive intrinsic properties to understand how *FMR1* reduction might render these deep layer pyramidal neurons hyperexcitable. Neither input resistance, membrane time constant (tau) nor sag slope were changed in sh*FMR1*-treated neurons, but their action potential waveform half width was significantly decreased. Because the maximum rising slope is increased alongside no changes in the falling slope (**Fig. 3b, d and Extended Data Fig. 6**), this finding points toward a stronger sodium channel activation following spike initiation.

Consistent with the changes in resting potential and altered AP waveform, we find multiple voltage-gated sodium channel (VGSC) genes are significantly upregulated in sh*FMR1* deep layer excitatory neurons, including auxiliary subunits *SCN1B* and *SCN2B* (**Fig. 3e, Extended Data Fig. 9a-b**). These subunits are similarly upregulated in FXS patient excitatory neurons (*SCN1B*: log2FC 0.90, p = 0.091; *SCN2B*: 0.873, p = 0.088), but either unchanged or significantly downregulated in *Fmr1^-/y^* excitatory neurons (**Fig. 3e and Extended Data Fig. 9b**). Together, these results show *FMR1* reduction increases cellular excitability in human cortical pyramidal neurons, with corresponding transcriptomic changes shared with FXS patients.

### *FMR1*-deficient human cortex exhibits network hyperexcitability

Resting-state EEG studies of FXS patients are suggestive of baseline hyperexcitability ^48,49^ and 12-18% of patients experience seizures with almost half of patients showing epileptiform spikes on electroencephalography^50,51^. Thus, it is important to understand whether acute *FMR1* reduction in human cortex causes network-level hyperexcitability. We transduced neurons with AAVrg-hSyn-GCaMP7s along with sh*FMR1* or shScr and performed 2p calcium imaging on 1-2 slice pairs from 6 patients at 8-10 days post-transduction **(Fig. 4a,b, Supplementary Table 1, Extended Data Fig. 10a)**. To unmask quiescent neurons that are inactive during the 15 min baseline recording, each recording was immediately followed by a 15 min exposure to a potent chemical stimulus to saturate neural activity (10 mM K^+^, 0.1 mM Mg^2+^, 0.1 mM 4-AP) and followed by a 15 min washout period **(Extended Data Fig. 10a)**. This design allowed us to both assess response to stimulation and confirm the viability of quiescent neurons under baseline conditions.

**Figure 4.**
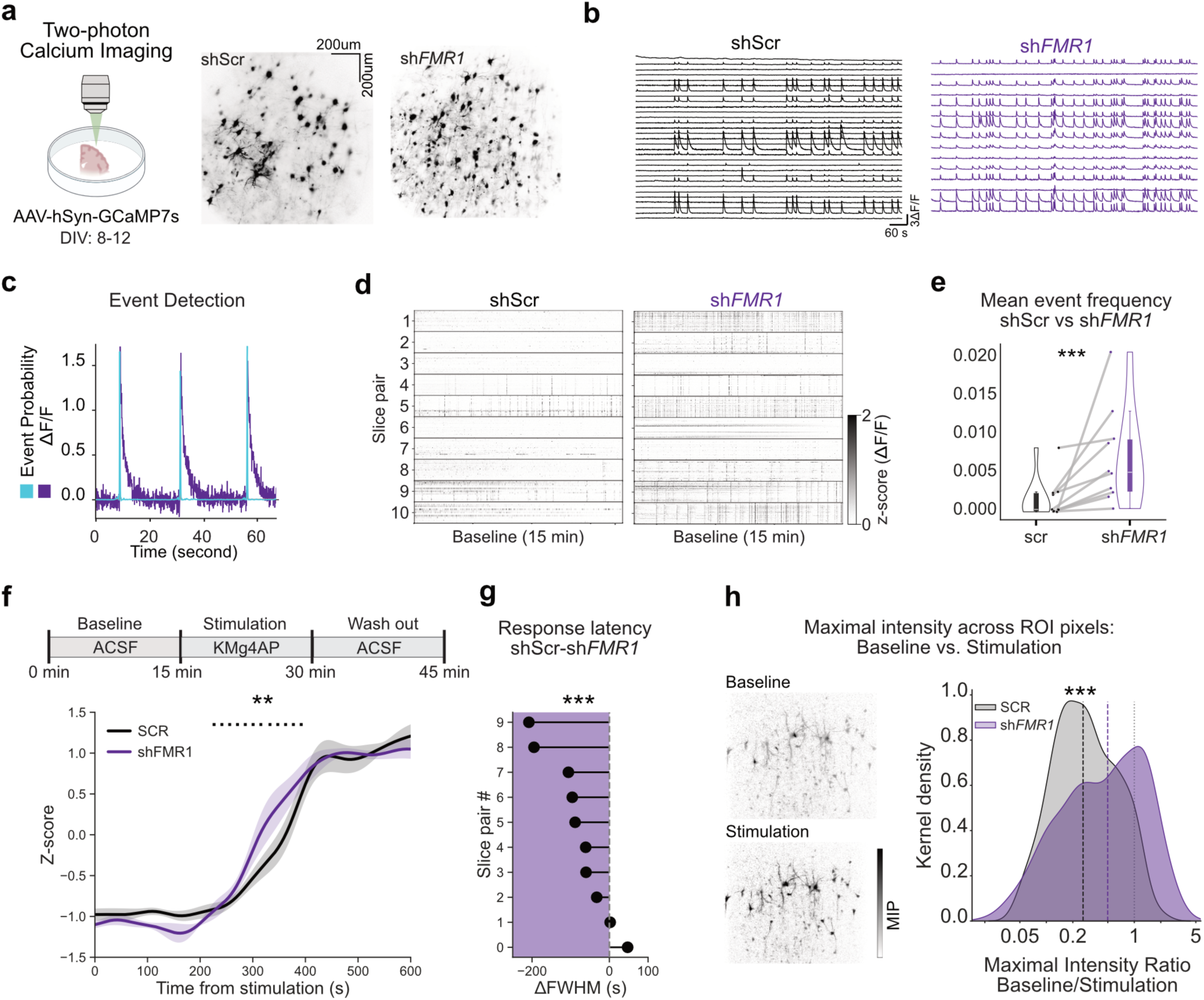
Reduced *FMR1* expression increases synchronized activity in human cortical microcircuits. **(a)** Representative 2P images showing the imaged neuronal population. **(b)** Example ΔF/F calcium traces from individual ROIs in shScr (left) and sh*FMR1* (right) slices**. (c)** Calcium transients from a representative neuron in a sh*FMR1* slice (purple) and the corresponding event-probability estimates generated using CASCADE (cyan). **(d)** Color-coded plots of calcium events in shScr (left) and sh*FMR1* (right) slices under baseline conditions. Each row represents a paired set of slices. Numbers indicate the number of detected neurons in each slice. (**e**) Mean event probability in shScr (black) and sh*FMR1* (purple) slices under baseline conditions. Each point represents the mean event probability across all detected neurons in one slice, and gray lines connect paired shScr and sh*FMR1* slices. Box plots show the median and interquartile range. \*\**P* < 0.01, generalized linear model with condition as the main factor. **(f)** Average low-pass-filtered calcium activity aligned to stimulation (10 mM K^+^, 0.1 mM Mg^2+^, 0.1 mM 4-AP) (KMg4AP) onset in sh*FMR1* (purple) and shScr (black) conditions. Shaded areas indicate s.e.m. (**g**) Differences in latency to half maximum (LHM) between conditions. \*\**P* < 0.01, generalized linear model with condition as the main factor. **(h)** Left, maximum-intensity projections (MIPs) during baseline (top) and stimulation (bottom) for an example slice. Right, kernel density distribution of the MIP ratio, calculated as baseline MIP divided by stimulation MIP and plotted on a logarithmic scale. \*\**P* < 0.01, generalized linear model with condition as the main factor.

Under baseline conditions, *FMR1*-deficient slices exhibit a significantly elevated rate of calcium events compared to paired shScr-treated slices from the same patient (**Fig. 4c-e and Extended Data Fig. 10b**). A linear mixed model considering all patient factors discussed above revealed a significant impact of brain region and donor age, similar to those seen in our analyses of cellular properties (**Supplementary Table 12)**. Upon application of the chemical stimulus to potently activate all neurons, including those that were quiescent at baseline, both groups of slices reached a robust maximum in activity, but sh*FMR1* slices reached this maximal response significantly faster **(Fig. 4f-g and Extended Data Fig. 10c).** Interestingly, an analysis of individual neuronal dynamics during stimulation and washout identified periods of slow oscillatory activity that were significantly different across sh*FMR1*- and shScr-treated slices (**Extended Data Fig. 10c**). To further understand the increased baseline activity of *FMR1*-deficient slices, we calculated the ratio of maximal activity at baseline versus stimulated conditions for each cell in sh*FMR1* and shScr slices. Shown as a ratio between Maximal Intensity Projection (MIP) images, sh*FMR1* cells exhibit a higher activity level at baseline that is much closer to the maximum of the chemical stimulus, and in many cases exceeding it **(Fig. 4h)**. Together, these results demonstrate that *FMR1* reduction dramatically increases the rate and magnitude of calcium activity, enhances sensitivity to stimulation and alters evoked dynamics in the human cortex, providing a clear and quantifiable signature of network changes.

## Conclusion

Collectively, our work represents the first multi-level analysis of *FMR1* reduction in human cortex that reveals alterations at the molecular, cellular, and network levels. It is interesting to note that many of the transcriptomic changes shared between *FMR1*-deficient slices and FXS patients are absent or discordant in the *Fmr1^-/y^* mouse cortex. This raises the possibility that there is a molecular impact of *FMR1* loss that is specific to human neurons. Given that the transcriptomic changes following acute *FMR1* reduction overlap with those of FXS patient cortex to a greater degree than those of ASD, bipolar disorder or Alzheimer’s disease cortex (**Fig. 2i-j**), it is likely that our functional results are more reflective of FMRP loss, than non-specific changes expected from working in the same species. Importantly, reducing *FMR1* acutely in cortex that has already developed, will undoubtedly not capture several FXS phenotypes thought to arise during critical period processes, including synapse maturation, experience-dependent plasticity and circuit stabilization^10,16,20^. Nonetheless, acute *FMR1* reduction in adult neurons can recapitulate molecular and circuit-level FXS phenotypes across models suggesting that many FXS symptoms may be due to the ongoing loss of FMRP^27,31,33^. Supporting this, FMRP replacement studies show that *FMR1* reintroduction resolves phenotypes even when administered in adult mouse brain^35,36^. The limitation of acute *FMR1* reduction only capturing the ongoing functions of FMRP may in fact prove to be a strength for therapeutic discovery, as phenotypes driven by the continued absence of FMRP are potentially more likely to be reversed with treatment. Thus, this approach to understanding genetic drivers of neurological disease in human cortex holds promise for identifying unique disease mechanisms that may not be represented in classic model systems.

## Materials and Methods (full Methods in Extended Data)

### Human brain slice culture

Human cortical tissue excised during epilepsy surgery was obtained with informed consent by the Repository Core for Neurological Disorders at BCH under IRB 09-02-0043. Patient information is listed in **Supplementary Table 1**. Tissue samples were excised from macroscopically normal cortex within the pathological context, adjacent to the lesioned area. Resected tissue was placed directly into ice-cold Slicing ACSF (saturated with 95% O2 / 5% CO2, with osmolarity of 305-315 mOsm and pH adjusted to 7.3) and transported to lab within 15 min. Tissue was trimmed, oriented and subjected to piotomy, after which 300 µm thick slices were prepared using a Leica VT 1200S vibratome. Slices were incubated in recovery solution then transferred to culture inserts and placed in the incubator overnight. Slices were transduced with AAV vectors after 24 hours.

### RNA extraction and RT-qPCR

Total RNA was extracted from cell suspension after FACS via TRIzol-chloroform extraction, and subsequent RNA clean-up of the resulting upper aqueous phase was performed with the RNA Clean & Concentrator™-5 kit (Zymo Research Cat# 11-325). cDNA was generated with the SuperScript™ IV VILO™ Master Mix (Invitrogen Cat# 11756050). For qPCR, the iTaq Universal SYBR Green Supermix (Bio-Rad Cat# 1725121) was used in combination with the Bio-Rad CFX Opus 96 System. For each sample, qPCR reactions were performed in triplicate for target gene (e.g. *FMR1*) and 2 reference genes (*ACTB* and *RPL13A*). The following primers were used: *FMR1* (Forward:GCAGCATGTGATGCAACTTACA; Reverse:CGCCTCTTTGGCACACATT); *ACTB* (Forward: CACCATTGGCAATGAGCGGTTC; Reverse: AGGTCTTTGCGGATGTCCACGT); *RPL13A* (Forward: CTCAAGGTGTTTGACGGCATCC; Reverse: TACTTCCAGCCAACCTCGTGAG). Primers were tested to have efficiency between 90% and 110%.

### Immunohistochemistry

Sections were subjected to antigen retrieval and passive delipidation (LifeCanvas Technologies), then immunostained using the following antibodies: mouse anti-FMRP (Developmental Studies Hybridoma Bank, clone 2F5-1; 1:500), rabbit anti-mCherry (Abcam, ab167453; 1:500), and chicken anti-NeuN (Abcam, ABN91; 1:500).

### Confocal Imaging

Sections were imaged for fluorescence using a Leica SP8 confocal. Stacks of images (250 μm, 2,5 μm z-steps) were acquired using a Leica HC Plain Apochromat 10X/0.40 CS2 objective with 1024 x 1024 μm pixel resolution or using 63X objective with zoom at 2x and 0.5 μm z-steps. Quantification analysis was performed using Imaris 10.0.0 and blind to treatment and genotype.

### Single-cell dissociation and Fluorescence Activated Cell Sorting (FACS)

Cortical slices transduced with AAV-shRNA for 7-10 days were dissociated using a papain-based protocol adapted from the Papain Dissociation System (Worthington Biochemical).

Single-cell suspensions were stained with DRAQ5 to label nucleated intact cells and DAPI was added just before sorting to exclude non-viable cells. Live neurons were defined as mCherry+ DRAQ5+ DAPI-singlets, with gates set using unstained controls, single-stained controls and non-transduced tissue. Sorted cells were collected into 1.5-mL tubes pre-coated with PBS lacking Ca^2+^/Mg^2+^ supplemented with 2% BSA/FBS and RNase inhibitor, using a small collection volume (∼20 - 30 µL) to maintain sufficient concentration for 10x loading.

### 10X genomics single-cell RNA-seq

Single-cell RNA-seq libraries were generated using the Chromium GEM-X Single Cell 3’ Kit v4 (10x Genomics; User Guide CG000731, Rev. B). For each sample (Scr and *FMR1* knockdown), live mCherry+ DRAQ5+ DAPI-neurons typically targeting 5000 cell recovery were loaded per channel of the Chromium X instrument for Gel Bead-in-Emulsion (GEM) generation. Libraries were purified with SPRI beads, and size distribution and integrity were assessed using an Agilent TapeStation. Library concentrations were quantified with a Qubit fluorometer. Indexed libraries were pooled and sequenced by Novogene (USA) on an Illumina NovaSeq-X plus platform (PE150-25B) using a standard 10x 3’ v4 read configuration (28 bp Read 1, 10 bp i7 index, 10bp i5 index, 90 bp Read 2). Scr and *FMR1* knockdown samples were processed and sequenced in parallel to minimize technical batch effects. Scr and *FMR1* samples from first patient sample (P1) were sequenced on a partial lane while others P2 and P3 were sequenced together on a full lane of NovaSeq-X plus.

### Single-cell RNA-seq processing

Fastq files were aligned to the human reference transcriptome GRCh38 and quantified using the count function from 10X Genomics Cell Ranger v8.0.1 with default parameters, yielding gene barcode and count matrices for each sample. Cell Ranger initially called a total of 25,953 cells combined in all samples. Raw matrices from Cell Ranger were then provided as input to CellBender ^52^ for background RNA removal and correction of ambient contamination. Pseudobulk count matrices were generated using Seurat’s (v 5.3.1) AggregateExpression function by summing raw RNA counts across cells within each condition × patient × merged cluster group, yielding one pseudobulk sample per donor per condition within each merged cell type. Differential expression was performed per merged cell type using only donors with paired sh*FMR1* and shScr samples (cell types with <3 pairs were excluded). To restrict analyses to robustly expressed genes, we applied donor-aware detection and low-count filters: genes were required to be detected (UMI>0) in ≥10% of cells in at least two paired donors and to have sufficient pseudobulk abundance (≥5 counts in ≥3 samples; genes with zero total counts were excluded).

### Cell type annotation

Cell type identification was done at level 1 using reference-based label transfer where scRNA-seq data (query) was mapped to a published human cortex reference atlas ^43^ using Seurat (v 5.3.1) ^53^. Both reference and query Seurat objects were independently processed and default assay for both was set to the RNA assay: counts were log-normalized (NormalizeData), highly variable genes were identified (FindVariableFeatures, 3,000 features), and expression values were scaled across variable features (ScaleData). Principal component analysis was computed for each dataset (RunPCA, 50 PCs). Cross-dataset anchors were identified with FindTransferAnchors using the reference PCA space (reference.reduction = “pca”, dims 1-30). Cell-type labels and sample-level covariates were transferred from the reference to the query using TransferData. Predicted labels and associated maximum prediction scores were added to the query metadata; for broad class labels, the full per-class prediction score matrix was retained. Finally, query cells were projected into the reference UMAP using MapQuery with the stored UMAP model (reduction.model = “umap”). Level 2 annotation was done using well established neuron-type marker gene expression (Extended Data Fig. 2a). Annotated populations include upper layer excitatory neurons: Exc_L2/3_LAMP5_ETV1, Exc_L2/3_CUX1_LAMP5, Exc_L2/4_CUX2_RORB, Exc_L4_RORB_CLSTN2, Exc_L4_RORB_PAX6; deep layer excitatory neurons Exc_L4/6_RORB_TSHZ2, Exc_L5/6_ETV1_FEZF2, Exc_L5/6_TLE4, Exc_L6_TLE4_CLSTN2, and inhibitory subclasses: Inh_CGE1_ADARB2-high, Inh_CGE2_ADARB2-low, Inh_SST_PV, Inh_VIP.

### Pseudobulk differential gene expression analysis

Pseudobulk count matrices were generated using Seurat’s (v 5.3.1) AggregateExpression function by summing raw RNA counts across cells within each condition × patient × merged cluster group, yielding one pseudobulk sample per donor per condition within each merged cell type. Differential expression was performed per merged cell type using only donors with paired sh*FMR1* and shScr samples (cell types with <3 pairs were excluded). To restrict analyses to robustly expressed genes, we applied donor-aware detection and low-count filters: genes were required to be detected (UMI>0) in ≥10% of cells in at least two paired donors and to have sufficient pseudobulk abundance (≥5 counts in ≥3 samples; genes with zero total counts were excluded). Differential expression testing was performed in DESeq2 (v 1.50.2) ^54^ using raw pseudobulk counts (rounded to integers) with a paired design (design: ∼ patient + condition). For each merged cell type, DESeq2 was run and contrasts were extracted as sh*FMR1* vs shScr (**Supplementary Table 3**). Results were reported as log2 fold change, Wald test statistic, nominal p-value, and Benjamini-Hochberg adjusted p-value (FDR).

For comparisons between datasets, pseudobulk analyses were performed on all excitatory and all inhibitory neurons of each dataset with the same gene filter and test (**Supplementary Table 6**). Counts were summed per donor (or animal) and condition, and genes were retained if they had ≥5 counts in at least k pseudobulk samples (k = number of donors or animals in the smallest group) and ≥5 counts in ≥10% of the samples of at least one condition; no cell-level detection filter was applied. Differential expression was tested with DESeq2 (Wald test); apeglm-shrunken log2 fold changes are reported in addition to the unshrunken estimates. For cross-species comparisons, mouse genes were mapped to human genes by one-to-one Ensembl orthologs.

### GO analysis

Gene Ontology (GO) analysis was performed on differential expression results using clusterProfiler (v 4.18.4) in R (v 4.5.0). For each DESeq2 (v 1.50.2) output table (CSV) containing gene, log2FoldChange, and a significance column (pvalue or padj). The background universe for each comparison was defined as the set of all detected gene symbols present in the corresponding DE table. Significant genes were selected using a fixed threshold on the pvalue < 0.01. GO enrichment was performed with enrichGO using org.Hs.eg.db (v 3.22.0) as the annotation database (keyType = “SYMBOL”), testing all GO ontologies (ont = “ALL”, encompassing BP, CC, and MF). Over-representation was evaluated relative to the per-comparison background universe and GO term p-values were adjusted using the Benjamini Hochberg (BH) method (pAdjustMethod = “BH”). For reporting, enrichment results were exported for each comparison as both TSV and Excel files, including GO term identifier and description, gene ratio, background ratio, Fold enrichment statistics, and contributing genes (**Supplementary Table 4**). In this workflow, the gene-level threshold for defining the input list of significant genes (pvalue < 0.01) is independent of the internal pvalueCutoff/qvalueCutoff settings passed to enrichGO, which were set permissively (both = 1) to return the full enriched-term table for downstream filtering and visualization.

### GSEA

Gene set enrichment analysis (GSEA) was performed using a pre-ranked approach implemented in the fgsea R package (v 1.36.2; fgseaMultilevel). For each comparison, genes were ranked by the DESeq2 Wald statistic, sorted from highest to lowest such that positive values corresponded to upregulation in the numerator condition and negative values to downregulation. Human gene symbols were converted to current HGNC symbols, and duplicate symbols were collapsed by retaining the entry with the largest absolute statistic. Gene sets were obtained from MSigDB via msigdbr (v 26.1.1; human GO collection C5), and terms were stratified into BP/CC/MF by MSigDB prefixes (GOBP_, GOCC_, GOMF_); each ontology was tested separately. Enrichment was computed using fgseaMultilevel with pathway size constraints (minSize = 10, maxSize = 500), eps = 0 and a fixed random seed. Results were reported as normalized enrichment score (NES), nominal p-value, and Benjamini Hochberg adjusted p-value (FDR), together with pathway size and leading-edge genes. For the slice subclusters (**Fig. 1g; Supplementary Table 5**), all genes tested in each subcluster were ranked. For comparisons across datasets (**Fig. 2c–e, Extended Data Fig. 4e–l; Supplementary Table 10**), GSEA was run on the excitatory and inhibitory analyses of all four datasets (human slice, FXS patient, Mouse-scRNA and Mouse-TRAP) using human C5 gene sets for every dataset (mouse genes via one-to-one orthologs), and the ranked lists were restricted to the genes tested in all eight analyses (8,845 genes), so that every dataset was tested on the same gene universe. Details of overlap analyses can be found in Extended Data.

Comparison with a second FXS patient cohort. Processed gene-level counts for GSE296077^59^ (post-mortem cortex, five FXS and five age-matched typically developing donors) were obtained from GEO. Genes detected at low abundance were removed before testing, and differential expression between FXS and typically developing donors was performed with DESeq2 using a single condition term; genes were ranked by the Wald statistic for enrichment analysis. Because this dataset profiles whole cortical tissue rather than sorted cell types, excitatory and inhibitory neurons were pooled into a single all-neuron pseudobulk profile per donor or animal for the human slice, the P5 *Fmr1^-/y^* scRNA-seq dataset and the adult *Fmr1^-/y^*TRAP-seq datasets (Camk2a-TRAP and Pvalb-TRAP combined), so that each comparator was matched to the cellular composition of the bulk reference. Overlap of differentially expressed genes (nominal *P* < 0.05 in both datasets) was assessed against size-matched random gene sets drawn from the genes tested in both datasets. For direction concordance, the concordance expected by chance was derived from each dataset’s proportion of positive-NES terms, and bars report the observed concordance of jointly significant terms (nominal *P* < 0.01) minus that expectation; confidence intervals and the two-sided *P* value for the human slice versus TRAP-seq difference were obtained from 5,000 bootstrap resamples of GO terms. Because bulk tissue contains glia, vasculature and blood in addition to neurons, differences in cellular composition between donor groups contribute to this comparison alongside cell-intrinsic differences in expression.

### Electrophysiology

Cortical slices transduced with AAV-shRNA for 7-10 days were removed from culture inserts and transferred to a recording chamber maintained at 31-32 °C and perfused with ACSF at a flow rate of 2 mL/min (ACSF: 120 mM sodium chloride, 2.5 mM potassium chloride, 1.25 mM sodium phosphate, 20 mM D-glucose, 26 mM sodium bicarbonate, 2 mM calcium chloride, 1 mM magnesium chloride, 1.3 mM sodium ascorbate, 3 mM myo-inositol and 3 mM sodium pyruvate, continuously bubbled with 95% O_2_ / 5% CO_2_). Patch-clamp recordings on deep L5/6 mCherry+ pyramidal neurons were performed with the aid of a fluorescence microscope (BX50WI, Olympus) with LED light source (pE-4000, CoolLED). Signals sampled at 20 KHz were passed through DigiData 1440A and amplified via a Multiclamp 700B amplifier, and recorded with pClamp software (version 10, Molecular Devices). Intrinsic properties were recorded in current-clamp following bridge balance-mediated series resistance compensation (Internal solution: 130 mM potassium gluconate, 10 mM potassium chloride, 10 mM Na phosphocreatine, 10 mM HEPES, 2 mM magnesium ATP, 0.3 mM sodium GTP, 0.2 mM EGTA and 1 mM magnesium chloride). One second hyperpolarizing and depolarizing step currents in 50 pA increments were injected to yield frequency-current (FI) curves. Resting potential, current threshold, sag currents and all active properties were extracted from the FI curve. Input resistance was calculated from the slope of the steady-state voltage response to hyper- and depolarizing step currents of smaller increments. Spike threshold was calculated as the voltage where time-differentiated voltage (dV/dt) exceeded 10 mV/ms. Sag slope was calculated via linear regression of the minimum hyperpolarizing sag voltage (within 250 ms of step current injection) as a function of the steady-state voltage (between 450 and 500 ms of step current injection). Sag decay time constant was calculated by exponential fit of the voltage response between minimum sag voltage and the end of the current step.

### Two-photon Calcium Imaging

Cortical slices were transduced with AAV-shRNA and AAV-hSyn-GCaMP7s ^51^ for 8-10 days prior to imaging to ensure robust expression. Imaging was focused on deep L5-6 cortex, identified by depth and laminar features. Two-photon calcium imaging was carried out on a Thorlabs Bergamo II microscope equipped with 920nm and 1064nm lasers (Spark) for GCaMP7s and images were acquired using a 16X Nikon CFI LWD Plan Fluorite 0.80 NA water-immersion objective. The field of view was 1024 × 1024 pixels, with a calibrated pixel size of 6.45 μm, and imaging was performed at 15 frames per second using Galvo-resonant scanning. During imaging, slices were continuously perfused with ACSF containing 120 mM sodium chloride, 3.5 mM potassium chloride, 1.25 mM sodium phosphate, 20 mM D-glucose, 26 mM sodium bicarbonate, 1 mM calcium chloride, 0.8 mM magnesium chloride, 1.3 mM sodium ascorbate, 3 mM myo-inositol and 3 mM sodium pyruvate. Temperature was maintained at 37°C using an automatic closed-loop temperature controller (Warner Instruments TC-344C), consistent with physiological ex vivo imaging conditions. Drug delivery was performed using a Minipuls 3 peristaltic pump (Gilson).

### Calcium imaging processing

Calcium imaging data were acquired as a RAW file in Thorlabs ImageLS software. Calcium movie data was processed using the Suite2p toolbox ^55^, available at https://www.github.com/cortex-lab/Suite2P. In brief, the Suite2p pipeline consists of registration, cell detection, region of interest (ROI) classification, neuropil correction and spike deconvolution. Movie frames are registered using 2D translation estimated by regularized phase correlation, subpixel interpolation and kriging. To detect ROIs (corresponding to cells), Suite2p clusters correlated pixels, using a low-dimensional decomposition of the data to accelerate processing. The number of ROIs is determined automatically by a threshold on pixel correlations. All detected ROIs were visually inspected and clustered into cells and not cells.

Because human cortical neurons are large, a single neuron occasionally produced multiple segmented ROIs in Suite2p. When a soma-containing ROI was present within the field of view, only that ROI was retained for analysis. In cases where the soma was not captured, we selected the ROI with the highest signal-to-noise ratio (SNR) as the representative ROI for a neuron.

Neuropil ROIs identified by Suite2p were excluded and not used in any subsequent analyses. For each ROI, we first computed the mean fluorescence signal *F(t)* by averaging pixel values within the ROI at each time point. To obtain a time-dependent baseline, we followed a sliding-window procedure adapted from previously described methods for calcium imaging ^56^. Briefly, the raw fluorescence trace was smoothed with a moving average (or Gaussian) filter with window length τ_1_ = 3 *seconds*. For each time point *t*, we then defined the baseline *F*_0_(*t*) as the minimum value of this smoothed trace within a preceding time window of length τ_2_ = 3 *seconds*, i.e. considering the interval [*t* - τ_2_, *t*]. The relative fluorescence change was computed as

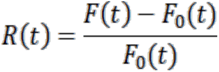

The resulting Δ*F/F(t)* traces were used for all subsequent event detection and population analyses (**Supplementary Table 12).**

### Calcium event detection

We used CASCADE, a deep-learning-based calcium deconvolution framework that estimates frame-wise spike probability from fluorescence traces, to infer significant calcium events ^57^. We used pre-trained model Global_EXC_30Hz_smoothing50ms_causalkernel. Time points with CASCADE-predicted spike probability greater than 0.5 were classified as significant calcium events.

### Latency to Half-Maximum (LHM) Measurement

To measure the latency of neuronal responses to stimulation, we computed the latency to half-maximum (LHM) during post-stimulation period. For each ROI calcium trace, we first low pass filtered ΔF/F signals with a 4^th^ order butterworth filter with a cutoff frequency of 0.01Hz to extract slow components of the signal. Next, we identified the maximum value of the filtered trace and computed the corresponding half-maximum threshold. We then detected the closest time point before the peak at which the signal crossed the half-maximum threshold. This timestamp represents the moment at which the transient first exceeded half of its maximum amplitude. Because the filtered trace captures slow envelope dynamics, this measure provides an estimate of the rise-time kinetics of calcium signal.

### Power Spectrum Analysis

To quantify low-frequency oscillatory activity in the calcium signal, the ΔF/F trace for each ROI was truncated to the analysis interval of interest. Unless otherwise specified, we analyzed the stimulation and wash epochs, which exhibited the most prominent rhythmic activity. The power spectral density (PSD) was computed using Welch’s method (Welch, 1967)^58^ with a 2-minute Hamming window and 50% overlap. PSD values were converted to decibels, PSD_dB_(*f*) = 10log_10_(PSD(*f*)), yielding a frequency-resolved spectrum that highlights dominant periodic components in the calcium activity.

### Maximal-intensity projection (MIP) analysis

Two-photon movies were motion-corrected and registered before maximal-intensity projection (MIP) analysis. Baseline and stimulation epochs were analyzed separately using the registered fluorescence movie, where x and y denote pixel position and t denotes imaging frame. For each pixel, baseline and stimulation MIPs were calculated as the maximum fluorescence intensity reached during the corresponding epoch:

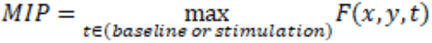

Thus, each MIP image represents the peak fluorescence value reached by each pixel during the corresponding imaging period. For visualization only, low-rank singular value decomposition reconstruction was used to reduce spatial noise, and a small median spatial filter was applied to reduce isolated single-pixel artifacts.

### AI statement

During the preparation of this work the authors used Claude (Anthropic, Version 2.31226.0) to check grammar and code. After using this tool, the authors reviewed and edited the content as needed and take full responsibility for the content of the publication.

## Acknowledgments

Experiments were performed in collaboration with the IDDRC core facilities at Boston Children’s Hospital (NIH P50 HD105351) including the Repository Core for Neurological Disorders, the Cellular Imaging Core, and the Flow Cytometry Core. Special thanks to all members of the NeuroBiorepository for assistance with tissue collection, Christopher A Walsh and Caroline M Dias to provide the FXS patient data, and to Gord Fishell for helpful comments on the manuscript.

## Funding

Wellcome Trust grant 219556/Z/19/Z (EKO)

FRAXA foundation grant (EKO)

Cell Discovery Network, Next-Gen Accelerator Grant 2025 (AS)

Developmental Neurology Institutional Training Grant, Boston Children’s Hospital, T32 NS007473 (MH)

Start-up and pilot funding TNC (EKO, JSF).

Esther A. & Joseph Klingenstein (JSF)

Simons Foundation (JSF)

CureSHANK (JSF)

Citizens United for Research in Epilepsy (JSF)

National Institutes of Health (NIH) (NS126725 to JSF).

Harvard Brain Science Initiative Bipolar Disorder Seed Grant (JSF)

Eagles Autism Foundation (EKO, JSF)

## Author contributions

Conceptualization: EKO, JSF

Methodology: AS, SA, LC, MH, SS, HL, BM, TPT, JY

Investigation: AS, SA, LC, MH, SS, HL, BM, TPT, JY, CMR, JT, EH, XX

Visualization: AS, MH, SA, EKO, JSF

Funding acquisition: EKO, JSF, AS

Project administration: AS, MH, SA, LC, CMR, JT, EH, EKO, JSF

Supervision: EKO, JSF, AS, MH, LC, SA

Writing – original draft: EKO, JSF, AS, MH, SA, JG

Writing – review & editing: all authors

## Competing interests

Authors declare that they have no competing interests.

## Data availability

Processed cell-level metadata, cluster annotations, pseudobulk differential expression results and pathway enrichment outputs are provided in Supplementary Tables 3-10, and electrophysiology statistics in Supplementary Table 11. Raw single-cell RNA-sequencing data will be deposited in dbGaP under controlled access to protect donor privacy; the accession number will be provided before publication. This study also analyzed the following published datasets: FXS patient and control cortex snRNA-seq (ref. 41; obtained from the authors); P5 *Fmr1*-KO mouse cortex scRNA-seq (ref. 38; GEO GSE147191); adult *Fmr1*-KO mouse Camk2a- and Pvalb-TRAP-seq (ref. 42; GEO GSE325852); autism spectrum disorder snRNA-seq (ref. 44; BioProject PRJNA434002); bipolar disorder snRNA-seq (ref. 45; GEO GSE306819); and Alzheimer’s disease snRNA-seq (SEA-AD, ref. 46; https://brain-map.org/consortia/sea-ad/our-data); and a second FXS patient and control cortex bulk RNA-seq dataset (ref. 43; GEO GSE296077). All other data supporting the findings of this study are available in the main text and Extended Data Information.

## Code availability

Single-cell RNA-seq data were processed and analysed with open-source software, including Cell Ranger, CellBender, Seurat, DESeq2, clusterProfiler and fgsea; calcium imaging data were analysed with Suite2p and CASCADE (see Methods). Custom R and Python scripts used for the analyses are available from the corresponding authors on request.

## Supplementary Information

Full Materials and Methods

Extended Data Figures 1-10

Supplementary Tables 1-12

Extended Data Movies 1-2

## Notes

### Competing Interest Statement

The authors have declared no competing interest.

### Summary of Updates

New results shown in Figures 2 and 3

