## Extended Data for "Acute *FMR1* reduction alters cellular and circuit properties in human cortex"

**The PDF file includes:**

Extended Data Figures
Materials and Methods
References 51-58

### Extended Data Figures

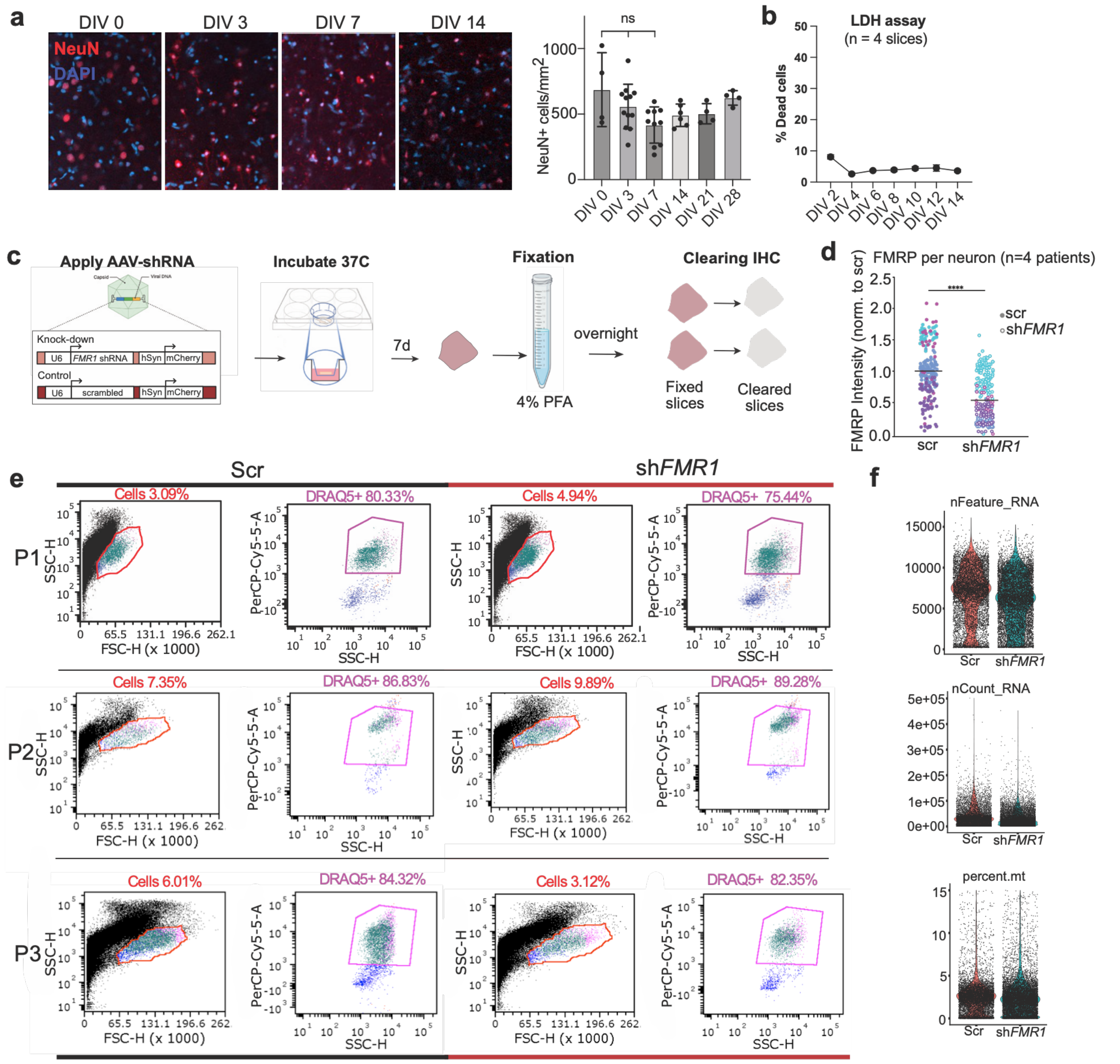


**Extended Data Fig. 1. Human cortical slices remain healthy over 4 weeks allowing significant reduction in *FMR1*.** **(a)** Representative immunohistochemistry (IHC) staining for neuron marker NeuN and DAPI at Days in vitro (DIV) 0, 3, 7 and 14. The number of NeuN+ cells per $mm^2$ was quantified in human cortical slices at DIV0-DIV28 to assess neuron viability. Quantification of multiple slices reveals an initial reduction (non-significant) followed by stable expression across 4 weeks. **(b)** Cell viability was assessed in human cortical slice cultures (N=4) by quantifying Lactate dehydrogenase (LDH) release in culture media across 14 DIV. This shows a stable low LDH level of < 10% across time in culture. **(c)** Schematic illustrating the experimental timeline for immunohistochemistry (IHC). Following viral transduction and 7 days of culture, cortical slices were fixed overnight and cleared during the IHC protocol. **(d)** Quantification revealed a significant reduction in FMRP intensity in shRNA-treated cortical slices compared to scrambled controls. Each color represents neurons from the same slice/patient. FMRP per neuron mean representation, n=4. Unpaired t-test $p<0.0001$. **(e)** Viable shRNA-expressing neurons were isolated from human cortical slices by FACs. Single-cell suspensions were stained with DRAQ5 to label nucleated intact cells, and DAPI to exclude non-viable cells, and run through a flow cytometer (BD FACS Aria). Debris and doublets were excluded based on forward and side scatter parameters and singlet gates (FSC-H vs

FSC-A, SSC-H vs SSC-A) (left panels; % of total events in the cell gate shown). Within this gate, cells were sorted by collecting the DAPI-/DRAQ5+ fraction (viable, nucleated population). This population represents an average of >75% of the total, indicating good viability post-dissociation for each condition. **(f)** Identification of number of unique genes (nFeature_RNA), total number of RNA molecules or UMIs (nCount_RNA) and percent of mitochondrial reads for shScr and sh*FMR1* populations is shown.

**a**

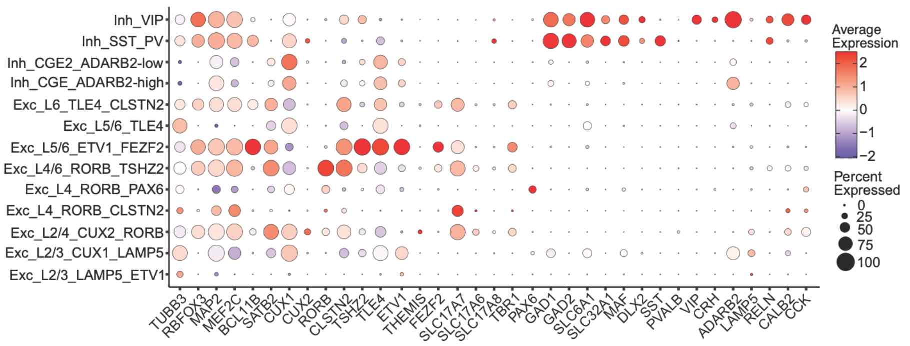


**b**

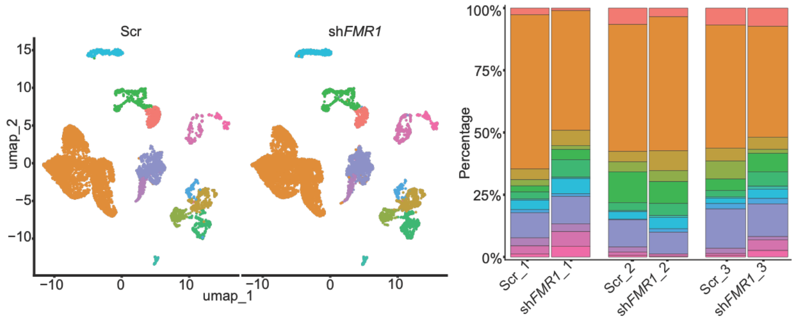


Number of cell in each cluster per condition

| Cell_types | Scr | sh*FMR1* |
|---|---|---|
| Exc_L2/3_LAMP5_ETV1 | 349 | 340 |
| Exc_L2/3_CUX1_LAMP5 | 4167 | 4702 |
| Exc_L2/4_CUX2_RORB | 339 | 590 |
| Exc_L4_RORB_CLSTN2 | 320 | 209 |
| Exc_L4_RORB_PAX6 | 358 | 592 |
| Exc_L4/6_RORB_TSHZ2 | 206 | 611 |
| Exc_L5/6_ETV1_FEZF2 | 38 | 73 |
| Exc_L5/6_TLE4 | 232 | 498 |
| Exc_L6_TLE4_CLSTN2 | 103 | 144 |
| Inh_CGE_ADARB2-high | 900 | 1098 |
| Inh_CGE2_ADARB2-low | 197 | 196 |
| Inh_SST_PV | 156 | 433 |
| Inh_VIP | 73 | 297 |

**Extended Data Fig. 2. Cell type identification for scRNA-seq. (a)** Dot plot shows canonical marker gene expression in identified neuron types from combined shScr and sh*FMR1* populations. **(b)** Left: UMAP plot shows identified clusters in shScr and sh*FMR1* populations separately. cell-type composition (middle) across excitatory and inhibitory classes show cellular subtypes present in both shScr and sh*FMR1*-expressing neurons from all 6 samples (N=3 patients). Right: Number of cells per cluster used in shScr and sh*FMR1* conditions in downstream analyses.

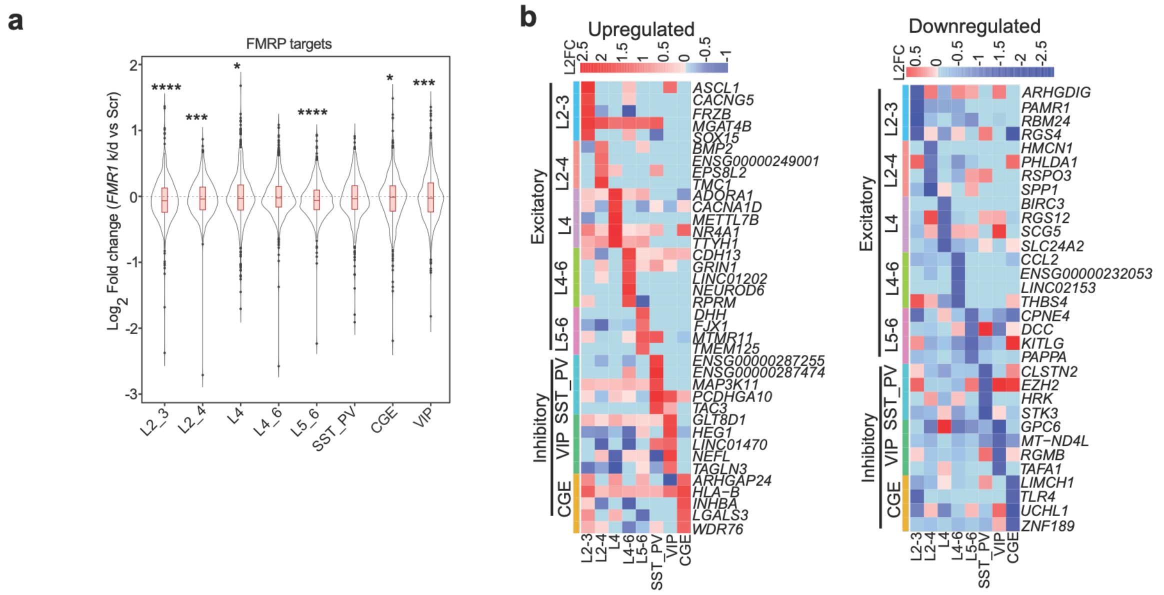


**Extended Data Fig. 3. Expression if FMRP target and differentially enriched genes in neuronal cell-types with *FMR1* reduction. (a)** FMRP target mRNAs are reduced in *FMR1*-deficient neurons. FMRP target mRNAs were identified from Dias et al., (2023) and expression quantified in neuron subtypes in sh*FMR1*-expressing neurons. Similar to what is seen in the *Fmr1*-KO mouse, our results show a downregulation in the mRNA targets of FMRP in almost all neuron types (violin plots; log2 fold-change, sh*FMR1* vs Scr; significant as indicated). Wilcoxon rank sum test, Exc_L2_3 (*P* =4.79075E-09), Exc_L2_4 (*P*=0.000326158), Exc_L4 (*P*=0.012145793), Exc_L4_6 (*P*=0.366251337), Exc_L5_6 (*P*=2.18777E-12), Inh_SST_PV, (*P*=0.433044542), Inh_CGE (*P*=0.014322204), Inh_VIP (*P*=6.71109E-05). **(b)** Heatmaps of differential gene expression ($Log_2$fold-change, sh*FMR1* vs shScr) show the most significantly upregulated and downregulated transcripts across excitatory and inhibitory subclasses.

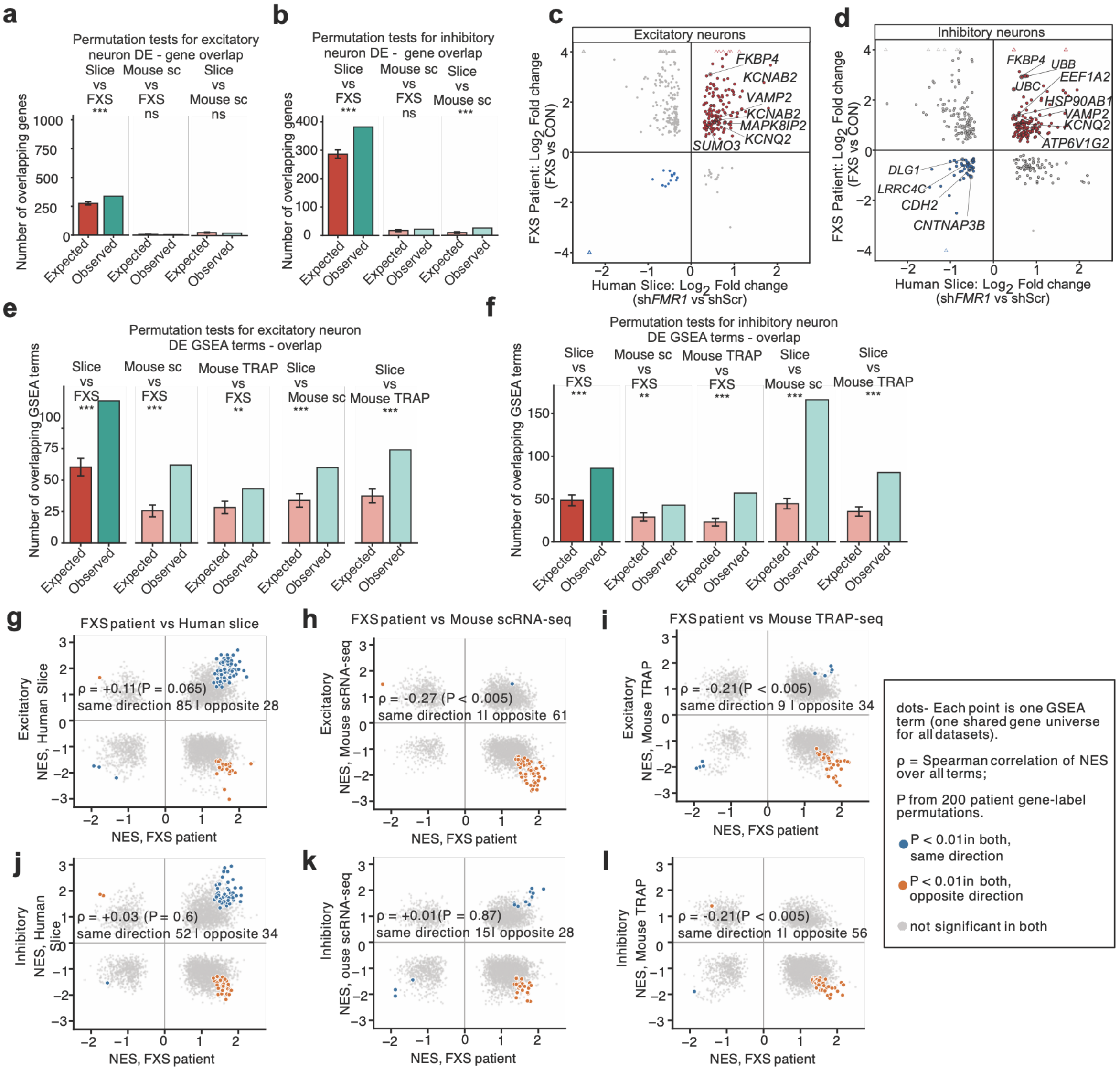


**Extended Data Fig. 4.** Gene and GSEA-term overlap and direction concordance between FXS patients, human slice and *Fmr1*-KO mouse datasets. Gene and GSEA-term overlap between FXS patient, human slice and *Fmr1*-KO mouse differential-expression datasets (same analyses as Fig. 2). **(a, b)** Gene overlap (DEGs, nominal $P < 0.05$ in both datasets) between human slice and FXS patients, P5 mouse scRNA-seq and FXS patients, and human slice and P5 mouse in excitatory **(a)** and inhibitory **(b)** neurons. Expected, mean ± s.d. of the hypergeometric distribution among genes tested in both datasets (matched by human Ensembl ID; mouse via 1:1 orthologs); *P*, permutation test (10,000 draws). Excitatory: slice vs FXS, N = 18,456, $n_{Slice}$ = 4,213, $n_{FXS}$ = 1,206, observed 338 vs expected 275 (1.23-fold), $P < 0.0001$; mouse vs FXS, N = 10,557, $n_{Mouse}$ = 90, $n_{FXS}$ = 769, 4 vs 6.6, $P$ = 0.90; slice vs mouse, N = 10,754, $n_{Mouse}$ = 95, $n_{Slice}$ = 2,897, 21 vs 25.6, $P$ = 0.88. Inhibitory: slice vs FXS, N = 16,533, $n_{Slice}$ = 1,731, $n_{FXS}$ = 2,736, 382 vs 286 (1.33-fold), $P < 0.0001$; mouse vs FXS, N = 8,987, $n_{Mouse}$ = 89, $n_{FXS}$ = 1,750, 22 vs 17.3, $P$ = 0.13; slice vs mouse, N = 8,973, $n_{Mouse}$ = 89, $n_{Slice}$ = 988, 26 vs 9.8 (2.65-fold), $P < 0.0001$. (c, d) Log2 fold change in the human slice (sh*FMR1* vs shScr; x axis) and FXS patients (FXS vs control; y axis) of genes with $P < 0.05$ in both datasets in excitatory **(c)** and inhibitory **(d)** neurons. Red, up in both; blue, down in both; grey, opposite direction; triangles, genes beyond the axis limits drawn at the edge; selected genes are labelled. **(e, f)** GSEA-term overlap (GO terms with nominal $P < 0.01$ in both datasets) for slice vs FXS, P5 mouse vs FXS, TRAP mouse vs FXS, slice vs P5 mouse and slice vs TRAP mouse in excitatory **(e)** and inhibitory **(f)** neurons. Expected, mean ± s.d. of the hypergeometric distribution among terms tested in both; *P*, one-sided Fisher's exact test. Excitatory, observed vs expected: 113 vs 60.3, 62 vs 25.5, 43 vs 28.2 ($P$ = 0.003), 60 vs 33.9 and 74 vs 37.4; inhibitory: 86 vs 48.6, 43 vs 29.1 ($P$ = 0.005), 57 vs 23.2, 166 vs 44.6 and

81 vs 35.6 (other $P < 0.0001$). (g–l) NES of GSEA terms in FXS patients (x axis) versus the human slice **(g, j),** P5 mouse scRNA-seq (h, k) and adult mouse TRAP-seq **(i, l)** in excitatory **(g-i)** and inhibitory **(j-l)** neurons. Each point is one GO term (one gene universe shared by all datasets). Blue, $P < 0.01$ in both, same direction; orange, $P < 0.01$ in both, opposite direction; grey, not significant in both. ρ, Spearman correlation of NES over all terms; *P* from 200 permutations of FXS patient gene labels (minimum $P = 0.005$). Numbers, jointly significant terms changed in the same or opposite direction. *$P < 0.05$, **$P < 0.01$, ***$P < 0.001$; ns, not significant.

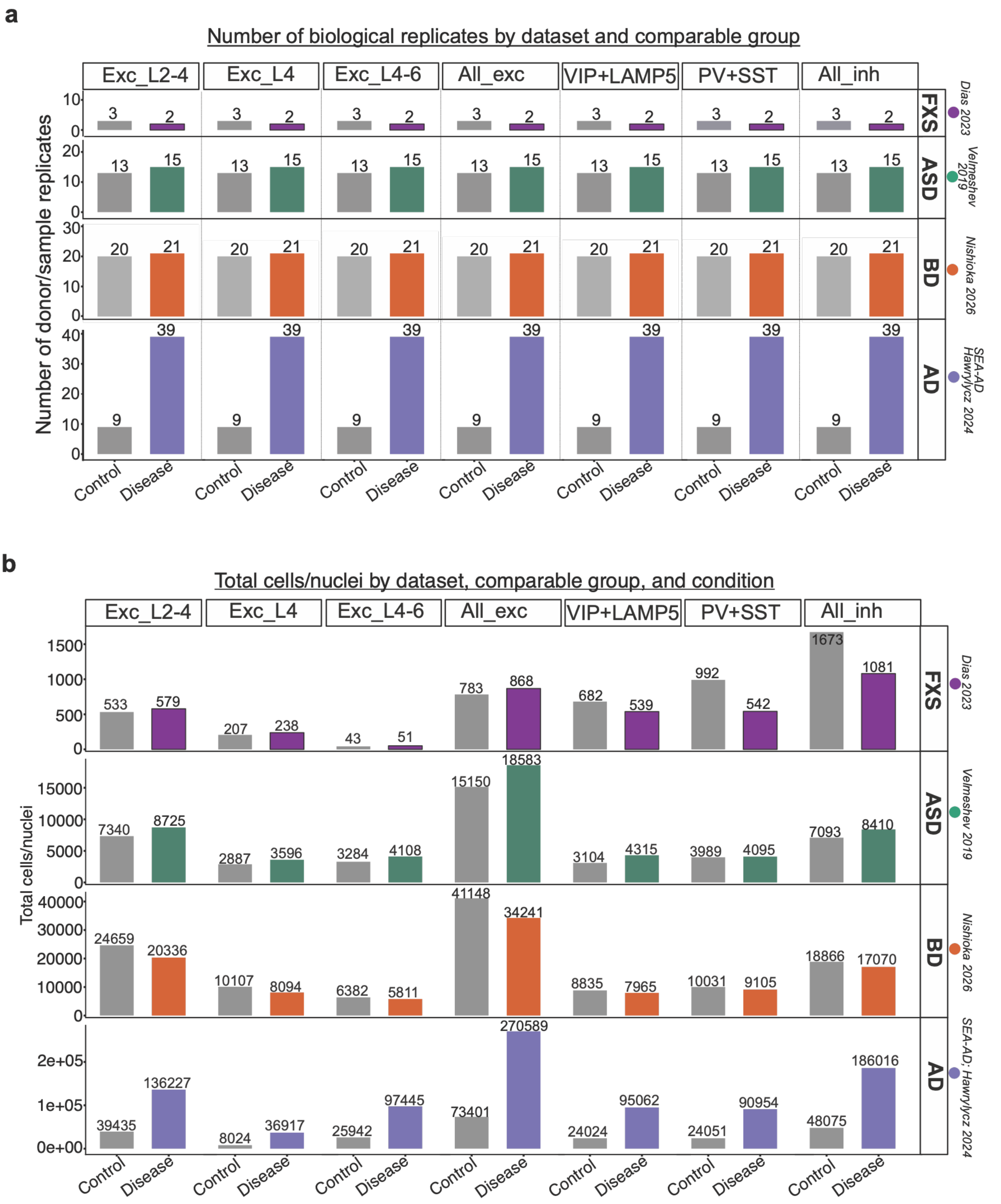


**Extended Data Fig. 5. Sample metrics and cell counts across datasets and cell-type groups. (a)** Number of biological replicates by dataset and comparable group. Bar plots showing the number of individual donor or sample replicates (y-axis) categorized by condition (Control vs. Disease; x-axis) across four distinct datasets (rows): Fragile X Syndrome (FXS, Dias 2023), Autism Spectrum Disorder (ASD, Velmeshev 2019), Bipolar Disorder (BD, Nishioka 2026), and Alzheimer's Disease (AD, Hawrylycz 2024). Metrics are broken down by major neuronal cell-type groups (columns), including excitatory layers (Exc_L2-4, Exc_L4, Exc_L4-6, and pooled All_exc) and inhibitory subtypes (VIP+LAMP5, PV+SST, and pooled All_inh). Exact sample sizes (n) are indicated above each bar.

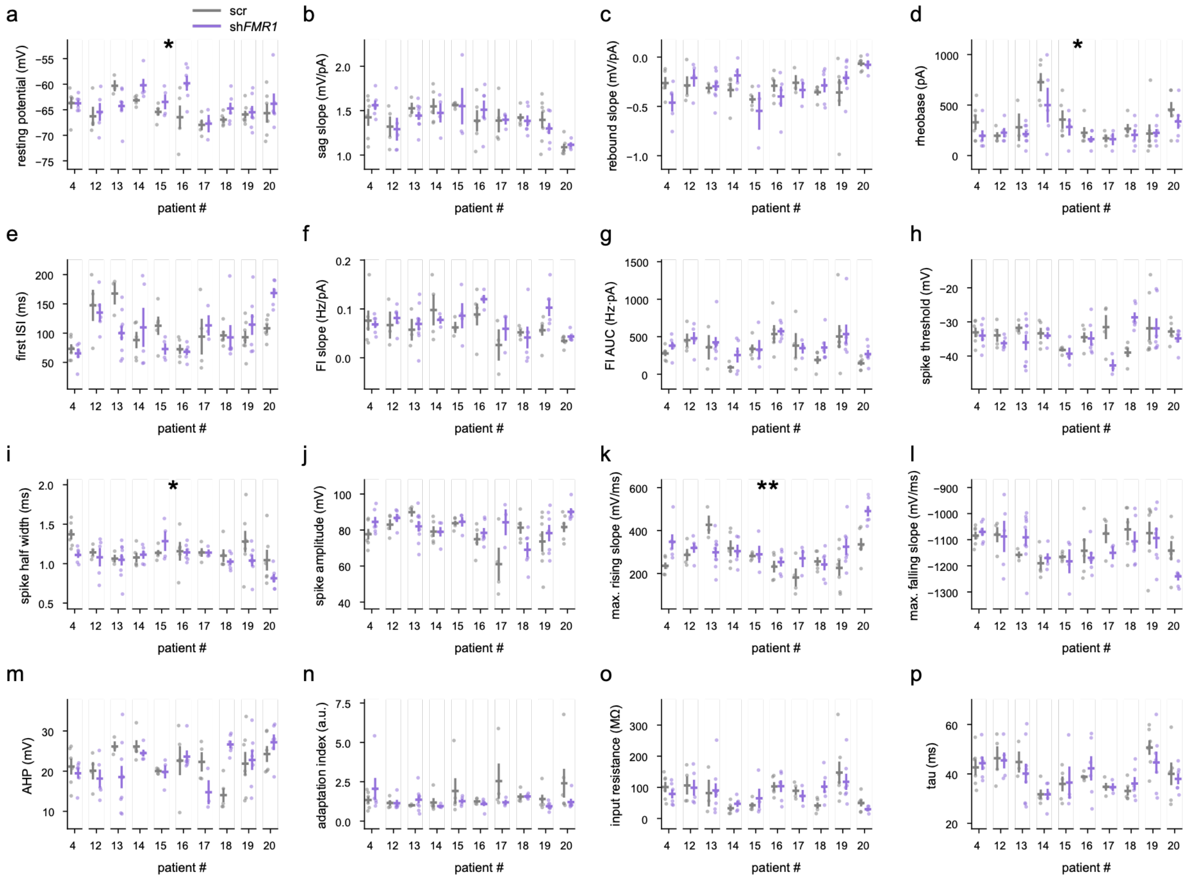

**Extended Data Fig. 6.** Patient-by-patient electrophysiological metrics of deep layer pyramidal neurons across scr- and sh*FMR1*-treated slices. **(a-p)** Scatter plots of each recorded neuron across patients with mean and SEM. * $P < 0.05$, ** $P < 0.01$, linear mixed-effects model with shRNA as main factor.

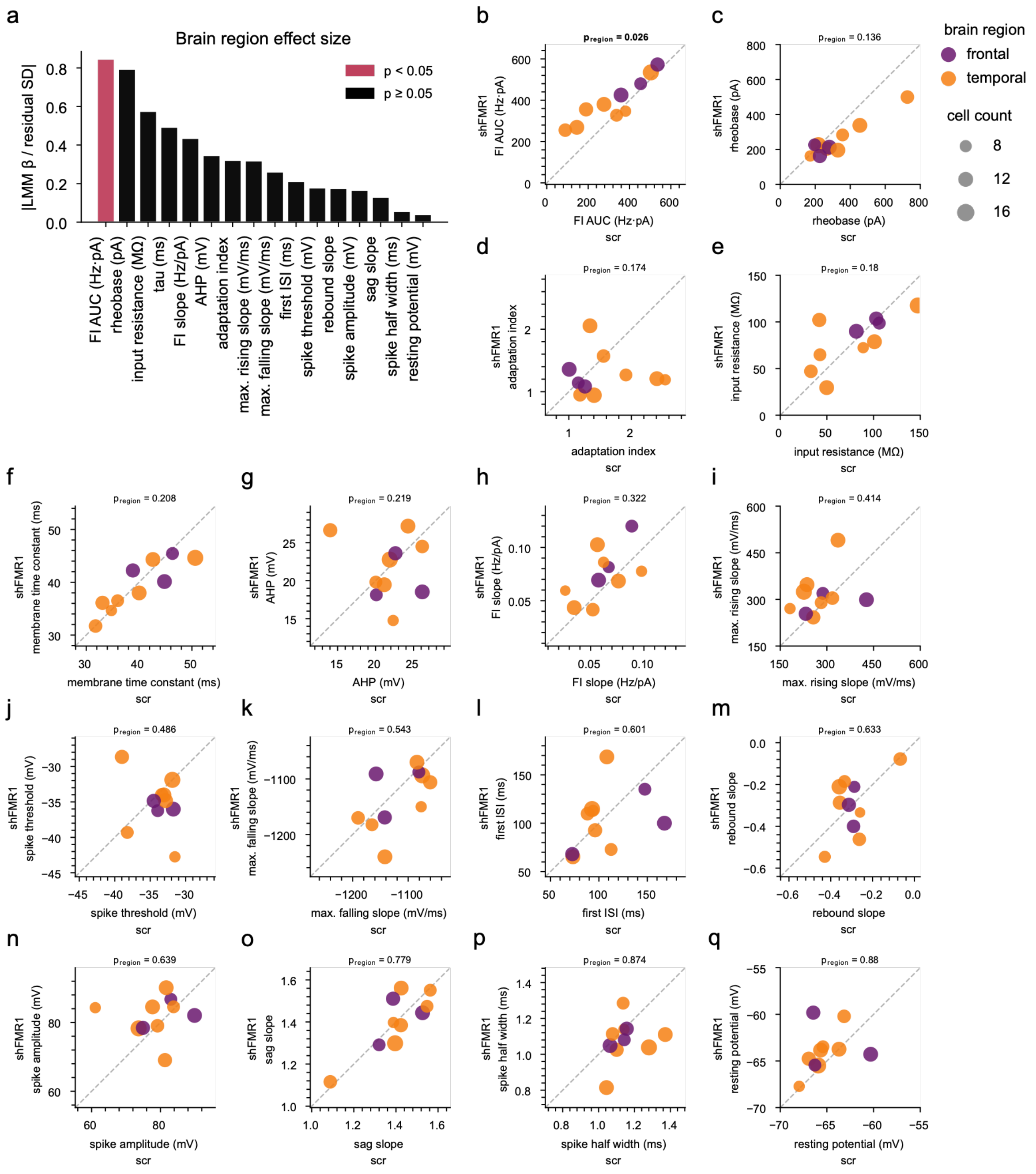


**Extended Data Fig. 7. Effect of brain region on deep layer pyramidal neuron excitability**. **(a)** Rank-ordered linear mixed-effects model (GLMM, see *Methods & Materials*) brain region effect sizes for each quantified electrophysiological metric. **(b-q)** Scatterplots for each electrophysiological metric with each dot representing each patient, positioned at the across-neuron average for scr- (*x*-axis) and shFMR1-treated (*y*-axis) neurons, size-weighted by total neurons recorded for that patient and colored by age. Scatterplots are rank-ordered by ascending *p*-value.

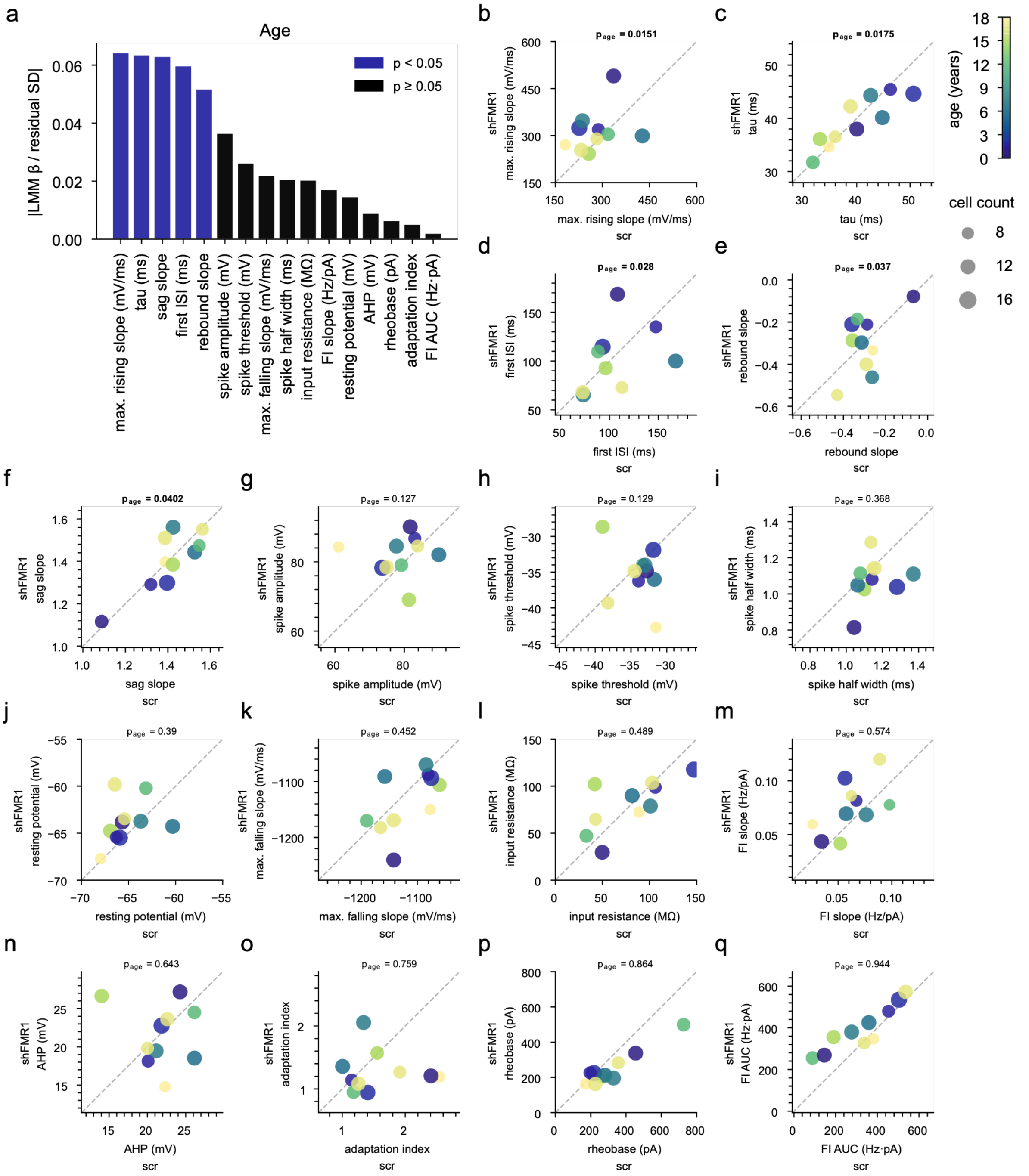


**Extended Data Fig. 8. Effect of age on deep layer pyramidal neuron excitability**. **(a)** Rank-ordered linear mixed-effects model (GLMM, see *Methods & Materials*) age effect sizes for each quantified electrophysiological metric. Effect size values are quantified per year. **(b-q)** Scatterplots for each electrophysiological metric with each dot representing each patient, positioned at the across-neuron average for scr- (*x*-axis) and shFMR1-treated (*y*-axis) neurons, size-weighted by total neurons recorded for that patient and colored by brain region. Scatterplots are rank-ordered by ascending *p*-value.

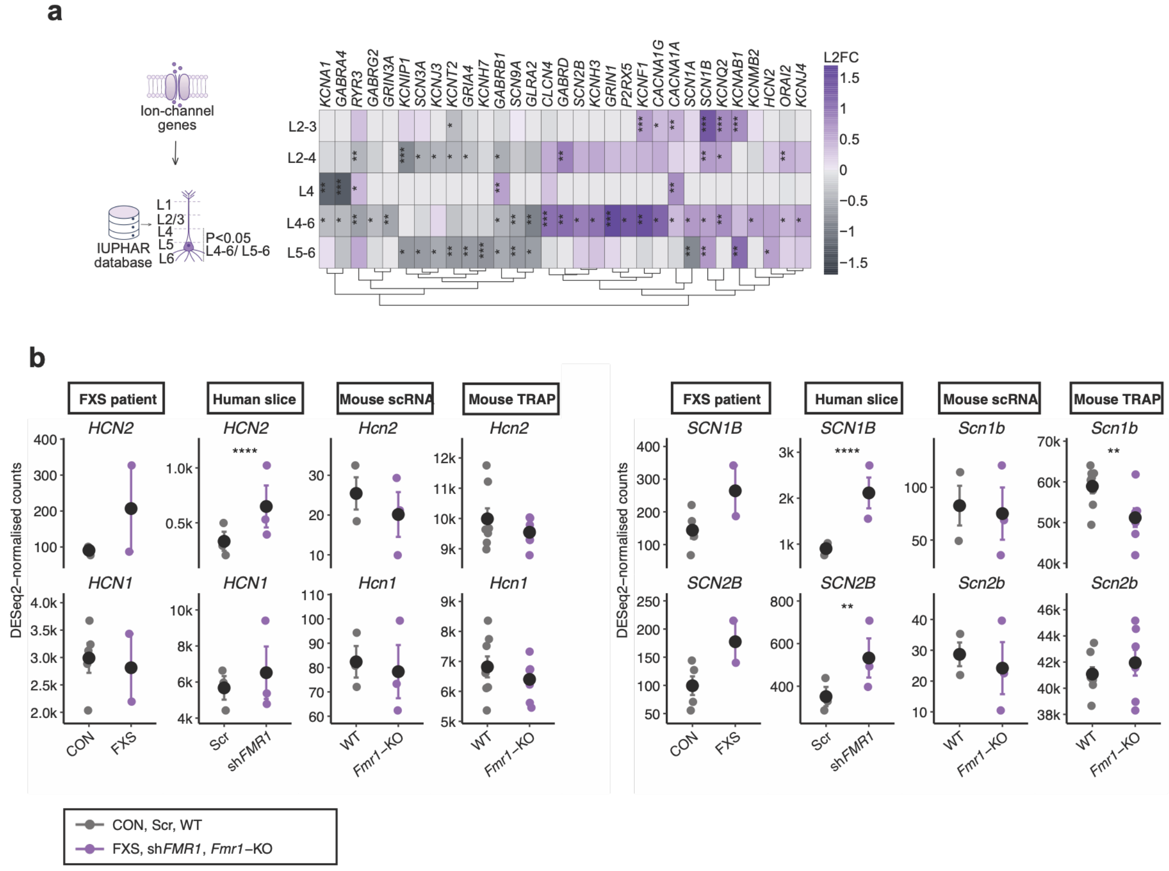


**Extended Data Fig. 9. Transcripts encoding ion channel subunits in the FXS patient, human slice and *Fmr1-KO* mouse model**. **(a)** Heatmap shows the differential expression of transcripts encoding ion channels in the excitatory neuron populations from each layer. Only significant (P<0.05) in deep layer L4-6 and L5-6 are selected. DESeq2 Wald test *P < 0.05, **P < 0.01., ***P < 0.001. **(b)** Deseq2 normalised expression of significantly altered ion channel subunit genes in excitatory neuron subclasses was compared between FXS patients, the *FMR1*-deficient Human model (sh*FMR1*), and the *Fmr1-KO* mouse model. For comparison across datasets, the collapsed mean log-normalised expression was plotted within the excitatory subclass. FXS patients show alterations in HCN channels and sodium channel subunits subunits that are recapitulated in the Human slice but not the *Fmr1-KO* mouse model * (P<0.05), ** (P<0.01), *** (P<0.001), **** (P<0.0001).

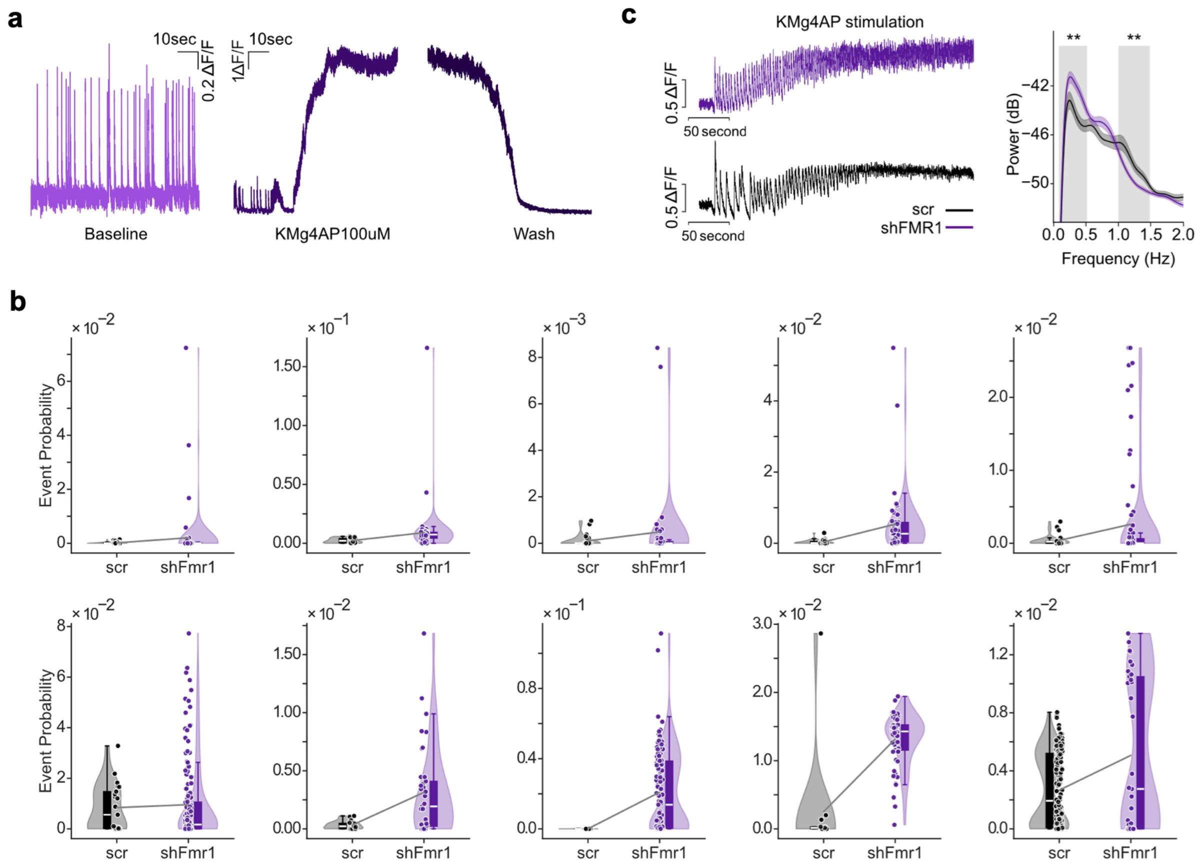


**Extended Data Fig. 10. Reduced *FMR1* expression increases calcium activity in neurons. (a)** Activity recorded at baseline (inset) and during stimulation is shown, with an experimental timeline of baseline activity in ACSF (15 min), stimulation with KMg4AP (100 µM; 15 min), followed by washout in ACSF (15 min). **(b)** Distribution of event probability across slice pairs. Each dot represents an ROI. Lines connect the average event probabilities of paired slices from the same donor (black: Scr; purple: sh*FMR1*). Each plot corresponds to a separate slice. **(c)** Left, example raw calcium traces during KMg4AP stimulation. Right, power spectra (mean ± 95% bootstrap CI) during KMg4AP and wash periods in sh*FMR1* (purple) and SCR (black) conditions. Gray regions denote frequency ranges with significant condition differences. **P < 0.01, two-tailed cluster-level permutation test, 5,000 permutations.

### Materials and Methods

#### Human brain slice culture

Human cortical tissue excised during epilepsy surgery was obtained with informed consent by the Repository Core for Neurological Disorders at BCH. Patient characteristics for the samples used in this study are listed in Supplementary Table 1. Tissue samples were excised from macroscopically normal cortex within the pathological context, adjacent to the lesioned area.

Resected tissue was placed directly into ice-cold Slicing ACSF (80mM NaCl, 75 mM Sucrose, 2.5mM KCl, 1.25mM $NaH_2PO_4$, 26mM $NaHCO_3$, 10 mM D-glucose, 0.5mM $CaCl_2$, 7mM $MgCl_2$, 1.3mM sodium ascorbate, 3.0 mM sodium pyruvate, 1x Penicillin-Streptomycin (Thermo Fisher, 15140122), and 3U/mL Nystatin (Sigma-Aldrich, N1638), saturated with 95% O2/ 5% CO2, with osmolarity of 305-315 mOsm and pH adjusted to 7.3) and transported to lab within 15 min. Tissue was trimmed, oriented and subjected to piotomy, after which 300 µm thick slices were prepared using a Leica VT 1200S vibratome. Slices were transferred to tissue culture media and placed in the incubator overnight. Slices were transduced with AAV vectors after 24 hours.

#### AAV transduction

AAV stocks were aliquoted and snap-frozen in liquid nitrogen for single-use. AAV transduction was performed immediately following fresh media change at 24 hr. Specifically, an AAV master mix was prepared consisting of 5 µL of each AAV construct, diluted to the desired titer with pre-warmed M2, and pipetted on top of the brain slice such that the liquid remained as a droplet on top of the slice and did not run down the sides. The following constructs were used in this study:

| Construct | Supplier | Source | Serotype | Final Titer |
|---|---|---|---|---|
| AAVrg-hSYN-mCherry-U6-scrmb-shRNA | Vector Biolabs | This study | AAVrg | 2.1 * 10^12 GC/mL |
| AAVrg-hSYN-mCherry-U6-h*FMR1*-shRNA | Vector Biolabs | This study | AAVrg | 2.2 * 10^12 GC/mL |
| AAV/rg-hSyn1-tdTomato-U6-scrmb | Vector Biolabs | This study | AAVrg | 6.5 * 10^12 GC/mL |
| AAVrg-hSYN-tdTomato-U6-h*FMR1*-shRNA | Vector Biolabs | This study | AAVrg | 6.5 * 10^12 GC/mL |
| pGP-AAV-syn-jGCaMP7s-WPRE | Addgene viral prep # 104487-AAVrg, RRID:Addgene_104487 | [51]Dana *et al.* 2019, Douglas Kim & GENIE Project | AAVrg | 1.3 * 10^13 GC/mL |

#### Lactate dehydrogenase (LDH) release assay

Overall slice viability was monitored by measuring lactate dehydrogenase (LDH), a cytosolic enzyme released into culture media by damaged cells. Media samples were collected 2 days after each media change and immediately frozen at -80ºC. Following collection of the final time point, a maximum-release sample was obtained for each slice by incubation with 1% Triton X-100 in fresh media for 2 hours at 37ºC before media collection. LDH was assayed in technical triplicate using the colorimetric CytoScan™ LDH Cytotoxicity Assay kit (G-Biosciences Cat# 786-324) according to the kit's protocol. Absorbance was measured at 490 nm (LDH signal) and 680 nm (background) using a CLARIOstar® *Plus* microplate reader (RRID:SCR_026330). For analysis, the 680 nm absorbance value was subtracted from the 490 nm value, the mean of technical replicates was averaged, and subsequently the fresh media's value was subtracted from the value of each sample to account for background signal from just media. Values at each time point were then normalized to the maximum-release value for each slice.

**RNA extraction and RT-qPCR**

Total RNA was extracted from cell suspension after FACS via TRIzol-chloroform extraction, and subsequent RNA clean-up of the resulting upper aqueous phase was performed with the RNA Clean & Concentrator™-5 kit (Zymo Research Cat# 11-325). cDNA was generated with the SuperScript™ IV VILO™ Master Mix (Invitrogen Cat# 11756050). For qPCR, the iTaq Universal SYBR Green Supermix (Bio-Rad Cat# 1725121) was used in combination with the Bio-Rad CFX Opus 96 System. For each sample, qPCR reactions were performed in triplicate for target gene (e.g. *FMR1*) and 2 reference genes (*ACTB* and *RPL13A*). The following primers were used: *FMR1* (Forward:GCAGCATGTGATGCAACTTACA; Reverse:CGCCTCTTTGGCACACATT); *ACTB* (Forward: CACCATTGGCAATGAGCGGTTC; Reverse: AGGTCTTTGCGGATGTCCACGT); *RPL13A* (Forward: CTCAAGGTGTTTGACGGCATCC; Reverse: TACTTCCAGCCAACCTCGTGAG). Primers were tested to have efficiency between 90% and 110%.

A modified Delta-Delta Ct method was used for analysis of qPCR data. Specifically, for each unique reaction defined by sample and gene being probed, technical replicate outliers where the coefficient of variation exceeds 2% were excluded, then the mean of remaining technical replicates was calculated. For each sample, Delta Ct (Dct) for each target gene (such as *FMR1*) was then calculated by subtracting the geometric mean of reference genes (*ACTB* and *RPL13A*) from the target gene. The Delta Dct (DDct) for each target gene was then calculated by subtracting the mean of Dct of all samples in the same run/plate from the Dct of each sample. Fold-difference is calculated as 2 to the power of negative DDct. Subsequently, the fold difference of each sample is divided by the mean fold-difference of the control group to enable comparison between experimental groups.

**Immunohistochemistry**

Sections were incubated in ethanol for 30 min, washed 2 x 10 min 0.3 M Phosphate Buffer Saline - 0.1% Tween 20 (PBST), and subjected to antigen retrieval in sodium citrate buffer (10mM, pH 6.0) at 85 °C for 30 min. Sections were cooled to room temperature for 5–10 min and washed 2 x 10 min 0.3 M PBST. Passive delipidation was performed overnight at room temperature using Delipidation Buffer (LifeCanvas Technologies). Sections were then washed 3 times (10-20min each) in 0.3 M PBST and blocked in 10% normal goat serum (NGS) in PBST for 2 h at room temperature. Primary antibody incubations were performed overnight at 4 °C in PBST containing 5% NGS using the following antibodies: mouse anti-FMRP (Developmental Studies Hybridoma Bank, clone 2F5-1; 1:500), rabbit anti-mCherry (Abcam, ab167453; 1:500), and chicken anti-NeuN (Abcam, ABN91; 1:500). Sections were washed 3 x 10 min PBST and incubated with secondary antibodies for 3 h at room temperature (all 1:500): goat anti-mouse Alexa Fluor 647, goat anti-chicken Alexa Fluor 488, and goat anti-rabbit Alexa Fluor 568 (Thermo Fisher). Sections were then washed 3 x 10 min PBST and post-fixed in 4% paraformaldehyde (PFA) for 30 min at room temperature, and index-matched using EasyIndex mounting medium (LifeCanvas Technologies) before mounting and imaging.

**Biocytin histochemistry**

Biocytin-filled cells were visualized through a streptavidin–biotin binding reaction. Following recordings, cortical slices were fixed in 4% paraformaldehyde for 24 hours at 4°C, washed 1× PBS, then transferred to 30% sucrose in PBS for 12–48 hours at 4°C. Sections were subjected to three freeze–thaw cycles, then blocked and permeabilized in block solution (PBS plus 10% NGS, 0.5% Triton X-100) for 2 hours at room temperature on an orbital shaker. Slices were then incubated overnight at 4°C on a rocker in blocking solution containing streptavidin conjugated to Alexa Fluor 647 (Invitrogen, S21374, 1:500) and DAPI (1:10,000 from a 10 µg/mL stock). After incubation, slices were washed 3 x 10min in PBS plus 0.5% Triton X-100, 2 x 10 min. PBS, then mounted onto slides using Fluoro-Gel mounting medium.

**Confocal Imaging**

Sections were imaged for fluorescence using a Leica SP8 confocal. Stacks of images (250 µm, 2,5 µm z-steps) were acquired using a Leica HC Plain Apochromat 10X/0.40 CS2 objective with 1024 x 1024 µm pixel resolution or using 63X objective for "per neuron quantification" with zoom at 2x and 0.5 µm z-steps. Quantification analysis was performed using Imaris 10.0.0 and blind to treatment and genotype. For "per neuron quantification", a mask was applied using NeuN signal and define individual neurons, from top to bottom. FMRP Sum intensity values were normalized to volume of neuron (with NeuN+ volume). Per slice a minimum of 15 neurons was counted.

**Single-cell dissociation**

Cortical slices transduced with AAV-shRNA for 7-10 days were dissociated using a papain-based protocol adapted from the Papain Dissociation System (Worthington Biochemical). Papain was dissolved to a final concentration of 20 U/mL in N-methyl-D-glucamine (NMDG)-based ACSF: 0.5mM $CaCl_2$, 25mM D-glucose, 20mM HEPES, 10mM $MgSO_4$, 1.25mM $NaH_2PO_4$, 3mM myo-inositol, 12mM N-acetylcysteine, 96mM NMDG, 2.5mM KCl, 25mM $NaHCO_3$, 5mM sodium L-ascorbate, 3mM sodium pyruvate, 0.01mM taurine and 2mM thiourea, adjusted to pH 7.4 with HCl, continuously bubbled with 95% $O_2$ / 5% $CO_2$. For each well of a 6-well plate, 800 µL papain solution and 40 µL of 2,000 U/mL DNase, were added and equilibrated for 15 min at 37ºC in a $CO_2$incubator. Immediately before adding tissue, 10 µL RNase inhibitor (Protector RNase Inhibitor, Roche) was added per well and gently swirled/mixed. Individual 300 µm slices were then transferred into the dissociation medium and incubated for 30 min at 37ºC, 5% $CO_2$, with gentle agitation every 5 min. After enzymatic digestion, plates were transferred onto ice and 200 µL albumin ovomucoid inhibitor solution was added to each well to quench papain activity. Slices were then gently triturated sequentially with fire-polished Pasteur pipettes of decreasing internal diameter (approximately 600 µm for 10 passes, 300 µm for 10 passes and 150 µm for 6 passes), avoiding air bubbles, to obtain a single-cell suspension. Cell suspensions were then transferred to 15mL falcon tubes pre-coated with ACSF containing 2% BSA/FBS and kept on ice. A 70 µm MACS smart strainer, pre-wetted with 2 mL ACSF + 2% BSA/FBS, was placed on top of each tube and 1 mL of dissociated cells was gently applied to the strainer. The strainer was subsequently washed with an additional 4 mL ACSF + 2% BSA/FBS. Cells were pelleted by centrifugation at 300g for 10 min at 4°C, the supernatant was carefully aspirated, and the pellet was resuspended in 500 µL ACSF with 2% BSA/FBS. The suspension was transferred to 1.5-mL tubes containing 10 µL RNase inhibitor and kept on ice. Cell number and viability were assessed using Trypan Blue exclusion and an automated cell counter.

**Fluorescence activated cell sorting (FACS)**
Single-cell suspensions were stained with DRAQ5 to label nucleated intact cells before leaving to the FACS facility and DAPI was added just before sorting to exclude non-viable cells. DRAQ5 (5 mM stock) was diluted 1:10 in PBS and then added to the cell suspension at a final 1:1,000 dilution; DAPI (1 mg/mL stock) was prepared similarly and added at a final 1:1,000 dilution immediately before sorting. Cells were kept on ice and protected from light during staining. Sorting was performed on a BD FACS Aria equipped with a 100-µm nozzle operated at low pressure. Debris and doublets were excluded based on forward and side scatter parameters and singlet gates (FSC-H vs FSC-A, SSC-H vs SSC-A). Neurons transduced with Scr or *FMR1* shRNA were identified by mCherry fluorescence from the AAV reporter. Live neurons were defined as mCherry+ DRAQ5+ DAPI- singlets*,* with gates set using unstained controls, single-stained controls and non-transduced tissue. Sorted cells were collected into 1.5-mL tubes pre-coated with PBS lacking $Ca^{2+}$/$Mg^{2+}$ supplemented with 2% BSA/FBS and RNase inhibitor, using a small collection volume (~20 - 30 µL) to maintain sufficient concentration for 10x loading.

**10X genomics single-cell RNA-seq library preparation and sequencing**
After sorting, cell number and viability were reassessed by Trypan Blue staining and manually counting cells using a Neubauer chamber, and the final cell concentration was adjusted according to 10x Genomics recommendations. Single-cell RNA-seq libraries were generated using the Chromium GEM-X Single Cell 3' Kit v4 (10x Genomics; User Guide CG000731, Rev. B). For each sample (Scr and *FMR1* knockdown), live mCherry+ DRAQ5+ DAPI- neurons typically targeting 5000 cell recovery were loaded per channel of the Chromium X instrument for Gel Bead-in-Emulsion (GEM) generation. Within each GEM, cells were lysed and polyadenylated transcripts were captured on barcoded gel beads, enabling incorporation of cell-specific barcodes and unique molecular identifiers (UMIs) during reverse transcription. After GEM generation and reverse transcription, emulsions were broken and cDNA was purified using magnetic beads. Full-length cDNA was amplified by PCR for 12 cycles, followed by enzymatic fragmentation, end-repair, A-tailing, adaptor ligation and sample index PCR for 11/13 cycles to generate final 3' gene expression libraries, following CG000731.

Libraries were purified with SPRI beads, and size distribution and integrity were assessed using an Agilent TapeStation. Library concentrations were quantified with a Qubit fluorometer. Indexed libraries were pooled and sequenced by Novogene (USA) on an Illumina NovaSeq-X plus platform (PE150-25B) using a standard 10x 3' v4 read configuration (28 bp Read 1, 10 bp i7 index, 10bp i5 index, 90 bp Read 2). Scr and *FMR1* knockdown samples were processed and sequenced in parallel to minimize technical batch effects. Scr and *FMR1* samples from first patient sample (P1) were sequenced on a partial lane while others P2 and P3 were sequenced together on a full lane of NovaSeq-X plus.

**Single-cell RNA-seq read alignment and gene expression quantification**
Fastq files were aligned to the human reference transcriptome GRCh38 and quantified using the count function from 10X Genomics Cell Ranger v8.0.1 with default parameters, yielding gene barcode and count matrices for each sample. Cell Ranger initially called a total of

25,953 cells combined in all samples. Raw matrices from Cell Ranger were then provided as input to CellBender [52] for background RNA removal and correction of ambient contamination. CellBender-filtered outputs were generated for each individual sample and used for downstream analysis. Following per-sample CellBender processing, datasets were integrated in Seurat using Reciprocal Principal Component Analysis (RPCA) based workflow. Briefly, each sample was log-normalized and 3000 variable features were selected, data were scaled, and principal component analysis (PCA) was performed. Samples were then integrated using the RPCA framework, followed by neighborhood graph construction, clustering, and UMAP dimensionality reduction. This initial integrated dataset contained 18,773 cells. To identify and remove putative doublets, scDblFinder was applied to the integrated object and predicted doublets were excluded. In parallel, mitochondrial quality filtering was applied and only cells with <15% mitochondrial (MT) reads were retained. After doublet removal and mitochondrial filtering, the dataset contained 17,441 cells. Clustering and UMAP were then recomputed on the filtered dataset, and downstream analyses were performed on these post-QC clusters. Cell types were assigned using a combination of reference mapping and canonical marker gene expression. Clusters lacking clear marker expression and/or containing fewer than 100 cells (combined for sh*FMR1* and shScr) were excluded. After stringent QC filtering, the final curated dataset comprised 17221 high quality cells, including 7438 shScr cells and 9783 sh*FMR1* cells.

Excitatory and inhibitory neurons across datasets. For comparisons between datasets, pseudobulk analyses were performed on all excitatory and all inhibitory neurons of each dataset with the same gene filter and test (**Supplementary Table 6**). Counts were summed per donor (or animal) and condition, and genes were retained if they had ≥5 counts in at least k pseudobulk samples (k = number of donors or animals in the smallest group) and ≥5 counts in ≥10% of the samples of at least one condition; no cell-level detection filter was applied. Differential expression was tested with DESeq2 (Wald test); apeglm-shrunken log2 fold changes are reported in addition to the unshrunken estimates. (i) Human slice: excitatory and inhibitory clusters of the 3 paired donors, design ~ patient + condition, sh*FMR1* vs shScr (27,953 excitatory and 20,774 inhibitory genes tested). (ii) FXS patient: single-nucleus RNA-seq of FXS (n = 2) and control (n = 5) cortex from Dias et al. (2023). Because donors had been processed with two versions of the gene annotation, gene symbols were converted to current HGNC symbols (limma alias2SymbolTable) and counts of synonymous symbols were merged before pseudobulk aggregation; genes detected only in donors processed with one annotation version were removed. Excitatory (L2/3-4, Neu NRGN, Neu L4 I and II, L5/6 I and II, Neu NTNG2) and inhibitory (Inh-SST, Inh-VIP, Inh-PVALB I and II, Inh-SV2C I and II) nuclei were summed per donor; control and FXS samples represented different individuals, therefore an unpaired design was used (design: ~ condition), FXS vs control (22,706 excitatory and 21,950 inhibitory genes tested). (iii) P5 *Fmr1*-KO mouse (Mouse-scRNA): single-cell RNA-seq of P5 cortex (Donnard et al., 2022; GSE147191). Mouse gene symbols were updated to current MGI symbols and merged, and excitatory neurons and interneurons were summed per animal (3 KO, 3 WT); design ~ condition, KO vs WT (11,666 excitatory and 9,790 inhibitory genes tested). (iv) Adult *Fmr1*-KO mouse (Mouse-TRAP): ribosome-associated mRNA from Camk2a-TRAP (excitatory; 7 KO, 8 WT) and Pvalb-TRAP (inhibitory; 10 KO, 10 WT) mice (Suresh et al., 2026; GSE325852). Immunoprecipitated counts from V1 and S1 were summed per animal; design ~ sex + genotype, KO vs WT (17,286 Camk2a and 17,517 Pvalb genes tested). For cross-species comparisons, mouse genes were mapped to human genes by one-to-one Ensembl orthologs.

**GO analysis**

Gene Ontology (GO) analysis was performed on differential expression results using clusterProfiler (v 4.18.4) in R (v 4.5.0). For each DESeq2 (v 1.50.2) output table (CSV) containing gene, log2FoldChange, and a significance column (pvalue or padj). The background universe for each comparison was defined as the set of all detected gene symbols present in the corresponding DE table. Significant genes were selected using a fixed threshold on the pvalue < 0.01. GO enrichment was performed with enrichGO using org.Hs.eg.db (v 3.22.0) as the annotation database (keyType = "SYMBOL"), testing all GO ontologies (ont = "ALL", encompassing BP, CC, and MF). Over-representation was evaluated relative to the per-comparison background universe, and GO term p-values were adjusted using the Benjamini Hochberg (BH) method (pAdjustMethod = "BH"). For reporting, enrichment results were exported for each comparison as both TSV and Excel files, including GO term identifier and description, gene ratio, background ratio, Fold enrichment statistics, and contributing genes (**Supplementary Table 4**). In this workflow, the gene-level threshold for defining the input list of significant genes (pvalue < 0.01) is independent of the internal pvalueCutoff/qvalueCutoff settings passed to enrichGO, which were set permissively (both = 1) to return the full enriched-term table for downstream filtering and visualization.

GO analysis of genes shared between the human slice and FXS patients (**Fig. 2a,b; Supplementary Table 9**) used genes with nominal P < 0.05 in both the sh*FMR1* slice and the FXS patient excitatory (or inhibitory) analyses, with all genes tested in both datasets (current HGNC symbols) as background. enrichGO was run as above (ont = "ALL", BH correction, gene-set size 10-500); terms with adjusted P < 0.05 and ≥5 genes were ranked by fold enrichment and the top 10 are shown.

**GSEA analysis**

Gene set enrichment analysis (GSEA) was performed using a pre-ranked approach implemented in the fgsea R package (v 1.36.2; fgseaMultilevel). For each comparison, genes were ranked by the DESeq2 Wald statistic, sorted from highest to lowest such that positive values corresponded to upregulation in the numerator condition and negative values to downregulation. Human gene symbols were converted to current HGNC symbols, and duplicate symbols were collapsed by retaining the entry with the largest absolute statistic. Gene sets were obtained from MSigDB via msigdbr (v 26.1.1; human GO collection C5), and terms were stratified into BP/CC/MF by MSigDB prefixes (GOBP_, GOCC_, GOMF_); each ontology was tested separately. Enrichment was computed using fgseaMultilevel with pathway size constraints (minSize = 10, maxSize = 500), eps = 0 and a fixed random seed. Results were reported as normalized enrichment score (NES), nominal p-value, and Benjamini Hochberg adjusted p-value (FDR), together with pathway size and leading-edge genes. For the slice subclusters (**Fig. 1g; Supplementary Table 5**), all genes tested in each subcluster were ranked. For comparisons across datasets (**Fig. 2c–e, Extended Data Fig. 4e–l; Supplementary Table 10**), GSEA was run on the excitatory and inhibitory analyses of all four datasets (human slice, FXS patient, Mouse-scRNA and Mouse-TRAP) using human C5 gene sets for every dataset (mouse genes via one-to-one orthologs), and the ranked lists were restricted to the genes tested in all eight analyses (8,845 genes), so that every dataset was tested on the same gene universe.

**Gene overlap analysis across datasets**

Overlap analysis between FXS, Slice and Mouse: Differential expression (DE) output tables were analyzed for - three datasets - FXS patient, human slice and Mouse-scRNA - separately for excitatory (Exc) and inhibitory (Inh) neuronal populations (Supplementary Table 6), and analyzed within cell class (Exc compared to Exc; Inh compared to Inh). Genes were matched by human Ensembl gene ID: human gene symbols were converted to Ensembl IDs (symbols mapping to a single ID only), and mouse genes were converted by one-to-one Ensembl orthologs; genes whose ID occurred more than once were excluded. Pairwise universes were defined as the intersection of tested genes between datasets (Exc / Inh: human slice and FXS patient, N = 18,456 / 16,533; Mouse-scRNA and FXS patient, N = 10,557 / 8,987; Mouse-scRNA and human slice, N = 10,754 / 8,973.

*Significance definition:* Within each dataset, genes were classified as significant using the unadjusted p-value threshold P < 0.05 (no directional filtering; both log2FC>0 and log2FC<0 included). For overlap testing within a given universe, significant sets were first restricted to genes present in that universe, so the effective list sizes (n_A, n_B) vary by universe (**Supplementary Table 7-8**).

Statistical testing of overlap enrichment: expected overlap under independence was (n_A × n_B)/N. Enrichment was assessed by a permutation test in which 10,000 overlaps were drawn from the hypergeometric distribution with the same N, n_A and n_B (equivalent to drawing random gene lists of size n_B from the universe), with P = (1 + number of draws ≥ observed overlap)/(10,000 + 1); one-sided Fisher's exact tests gave the same conclusions (Supplementary Table 7). For the three-way comparison (**Supplementary Table 8**), genes tested in the FXS patient, human slice and mouse datasets were classified as significant in all three, or in only two of the three datasets.

*Significance definition:* Within each dataset, genes were classified as significant using the unadjusted p-value threshold $P < 0.05$ (no directional filtering; both log2FC>0 and log2FC<0 included). For overlap testing within a given universe, significant sets were first restricted to genes present in that universe, so the effective list sizes (n_A, n_B) vary by universe (**Supplementary Table 7-8**).

*Statistical testing of overlap enrichment:* Overlap enrichment of significant genes was evaluated using a one-sided Fisher's exact test ("greater") on the 2×2 contingency table defined within the tested-gene universe. Expected overlap under independence was approximated as (n_A × n_B)/N. Effect sizes are reported as odds ratio (**Supplementary Table 7**).

Overlap analysis between Slice model and disease datasets: For comparisons between the human slice model and larger patient disease datasets, including ASD, bipolar disorder, and Alzheimer's disease, we performed an additional FXS-anchored resampling analysis to account for differences in DEG list size. Within each harmonized neuronal class, the number of FXS patient DEGs in the Slice-FXS pairwise universe was used as the matched disease-signature size. The observed Slice-FXS overlap was calculated directly. For ASD, bipolar disorder, and Alzheimer's disease, an FXS-matched number of genes was repeatedly sampled from the full pool of nominal disease DEGs ($p < 0.05$) within the corresponding Slice-disease pairwise universe, and overlap with the slice DE signature was recalculated for each selection. To estimate expected overlap, matched-size random gene sets were sampled from the full pairwise tested-gene universe, irrespective of disease DE status. Overlap was reported as the percentage of selected disease DE genes captured by the slice DE signature, ignoring direction of change.

**GSEA overlap analysis across datasets**

GSEA output tables (see GSEA analysis; one shared gene universe) were analyzed for the four datasets separately for excitatory (Exc) and inhibitory (Inh) neuronal populations (**Supplementary Table 10**). Gene sets were defined by the GSEA identifier and analyzed within cell class (Exc compared to Exc; Inh compared to Inh). Pairwise universes were defined as the intersection of gene sets tested in each dataset pair (Exc N = 4,956–4,959; Inh N = 4,959).

Significance definition: Within each dataset, a gene set was classified as significant using the unadjusted p-value threshold $P < 0.01$ (both NES > 0 and NES < 0 included). For overlap analyses within a given universe, significant sets were first restricted to the gene sets contained in that universe; thus, the effective set sizes (n_A, n_B) vary by universe.

Statistical testing of overlap enrichment: Overlap enrichment of significant gene sets was evaluated within each pairwise intersected universe using a one-sided Fisher's exact test ("greater") on the corresponding 2×2 contingency table. Expected overlap under independence was calculated as (n_A × n_B)/N, with s.d. from the hypergeometric distribution (Extended Data Fig. 4e,f).

**GSEA direction concordance and NES correlation**
For gene sets significant (P < 0.01) in both the FXS patient and a comparison dataset (Fig. 2d), we calculated the percentage with the same NES sign, with a 95% Wilson confidence interval. The concordance expected by chance was p_A·p_B + (1 − p_A)(1 − p_B), where p_A and p_B are the proportions of significant gene sets with positive NES in each dataset, and observed concordance was compared with this expectation by a two-sided binomial test. The Spearman correlation of NES across all gene sets (Extended Data Fig. 4g–l) was tested against 200 permutations in which the FXS patient gene ranks were shuffled and GSEA was re-run (two-sided; minimum P = 0.005). For **Fig. 2e**, gene sets significant (P < 0.01) in both the human slice and the FXS patient with the same NES sign were ranked by mean |NES|.

**Comparison with a second FXS patient cohort**
Processed gene-level counts for the bulk RNA-sequencing dataset GSE296077 (post-mortem cortex, five FXS and five age-matched typically developing donors) were obtained from GEO. Genes detected at low abundance were removed before testing, and differential expression between FXS and typically developing donors was performed with DESeq2 using a single condition term; genes were ranked by the Wald statistic for enrichment analysis, as above. Because this dataset profiles whole cortical tissue rather than sorted cell types, excitatory and inhibitory neurons were pooled into a single all-neuron pseudobulk profile per donor or animal for the human slice, the P5 *Fmr1*$^{-/y}$ scRNA-seq dataset and the adult *Fmr1*$^{-/y}$ TRAP-seq datasets (Camk2a-TRAP and Pvalb-TRAP combined), so that each comparator was matched to the cellular composition of the bulk reference. Overlap of differentially expressed genes (nominal $P < 0.05$ in both datasets) was assessed against size-matched random gene sets drawn from the genes tested in both datasets (Fig. 2f). For direction concordance (Fig. 2h), the concordance expected by chance was derived from each dataset's proportion of positive-NES terms, and bars report the observed concordance of jointly significant terms (nominal $P < 0.01$) minus that expectation; confidence intervals and the two-sided $P$ value for the human slice versus TRAP-seq difference were obtained from 5,000 bootstrap resamples of GO terms. Because bulk tissue contains glia, vasculature and blood in addition to neurons, differences in cellular composition between donor groups contribute to this comparison alongside cell-intrinsic differences in expression.

**Comparison with other neurological disorders**
To test whether the sh*FMR1* slice preferentially captures FXS-associated changes (**Fig. 2i,j**), cell types of the human slice and of single-nucleus datasets of FXS (Dias et al., 2023), autism spectrum disorder (ASD; Velmeshev et al., 2019), bipolar disorder (BD; Nishioka et al., 2026) and Alzheimer's disease (AD; Hawrylycz et al., 2024) were harmonized into seven comparable groups (L2-4, L4 and L4-6 excitatory neurons, all excitatory neurons, VIP+LAMP5 and PV+SST interneurons, and all inhibitory neurons), and pseudobulk DESeq2 analysis was performed per group and dataset (FXS patient as described above, by cell-type group; Extended Data Fig. 5). For each group and disease, the overlap was the percentage of disease DE genes (P < 0.05) that were also DE in the sh*FMR1* slice (P < 0.05), among genes tested in both datasets. To compare diseases with different numbers of DE genes, ASD, BD and AD DE gene lists were down-sampled to the number of FXS DE genes in the same group (1,000 random draws; mean reported). The background was estimated from 10,000 random gene sets of the same size drawn from the same universe (mean ± s.d.), P values were calculated against this background and corrected across the 28 tests (7 groups

× 4 diseases) with the Benjamini–Hochberg method. Relative overlap (heatmap) was expressed as a percentage of the FXS overlap in the same group.

**Ion-channel gene expression across datasets**
Ion channel gene list was used from IUPHAR database and HGNC symbols cobined. Log2 fold changes of ion-channel and receptor genes in excitatory neurons (Fig. 3e) were taken from the DESeq2 analyses described above (FXS patient, human slice, Mouse-scRNA and Camk2a Mouse-TRAP; mouse genes via one-to-one orthologs); asterisks show nominal DESeq2 Wald test P values.

All electrophysiological analyses and statistics were performed using custom Python scripts. Linear mixed-effects models were fit (**Supplementary Table 11**) to assess the effects of shRNA condition, sex and brain region on each electrophysiological metric using the 'statsmodels' framework in Python. Repeated neuron measurements within individual patients were accounted for by including patient labels as a random intercept.

**Two-photon Calcium Imaging**
Cortical slices were transduced with AAV-shRNA and AAV-hSyn-GCaMP7s [51] for 8-10 days prior to imaging to ensure robust expression. Imaging was focused on deep L5-6 cortex, identified by depth and laminar features. Two-photon calcium imaging was carried out on a Thorlabs Bergamo II microscope equipped with 920nm and 1064nm lasers (Spark) for GCaMP7s and images were acquired using a 16X Nikon CFI LWD Plan Fluorite 0.80 NA water-immersion objective. The field of view was 1024 × 1024 pixels, with a calibrated pixel

size of 6.45 µm, and imaging was performed at 15 frames per second using Galvo-resonant scanning. During imaging, slices were continuously perfused with ACSF containing 120 mM sodium chloride, 3.5 mM potassium chloride, 1.25 mM sodium phosphate, 20 mM D-glucose, 26 mM sodium bicarbonate, 1 mM calcium chloride, 0.8 mM magnesium chloride, 1.3 mM sodium ascorbate, 3 mM myo-inositol and 3 mM sodium pyruvate. Temperature was maintained at 37°C using an automatic closed-loop temperature controller (Warner Instruments TC-344C), consistent with physiological ex vivo imaging conditions. Drug delivery was performed using a Minipuls 3 peristaltic pump (Gilson). Each imaging session consisted of three continuous 15-minute conditions: 1) Baseline: ACSF only; 2) Stimulation: modified ACSF containing 10 mM KCl, 0.1 mM $MgCl_2$ and 100 µM 4-AP to elevate network excitability; 3) Wash: Return to ACSF. Transitions between conditions were performed using the peristaltic pump system.

**Calcium imaging processing**

Calcium imaging data were acquired as a RAW file in Thorlabs ImageLS software. Calcium movie data was processed using the Suite2p toolbox [55], available at https://www.github.com/cortex-lab/Suite2P. In brief, the Suite2p pipeline consists of registration, cell detection, region of interest (ROI) classification, neuropil correction and spike deconvolution. Movie frames are registered using 2D translation estimated by regularized phase correlation, subpixel interpolation and kriging. To detect ROIs (corresponding to cells), Suite2p clusters correlated pixels, using a low-dimensional decomposition of the data to accelerate processing. The number of ROIs is determined automatically by a threshold on pixel correlations. All detected ROIs were visually inspected and clustered into cells and not cells. Because human cortical neurons are large, a single neuron occasionally produced multiple segmented ROIs in Suite2p. When a soma-containing ROI was present within the field of view, only that ROI was retained for analysis. In cases where the soma was not captured, we selected the ROI with the highest signal-to-noise ratio (SNR) as the representative ROI for a neuron. Neuropil ROIs identified by Suite2p were excluded and not used in any subsequent analyses.

**Suite2p Parameters**

| Parameter | Value |
|---|---|
| Input format | raw |
| nchannels | 1 |
| Functional channel | 1 |
| Fs | 15 |
| roidetect | 1 |
| sparse_mode | 1 |

| | |
|---|---|
| Anatomical only | 0 |
| Neuropil_extract | 1 |

For each ROI, we first computed the mean fluorescence signal $F(t)$ by averaging pixel values within the ROI at each time point. To obtain a time-dependent baseline, we followed a sliding-window procedure adapted from previously described methods for calcium imaging [56]. Briefly, the raw fluorescence trace was smoothed with a moving average (or Gaussian) filter with window length $\tau_1 = 3\ seconds$. For each time point $t$, we then defined the baseline $F_0(t)$ as the minimum value of this smoothed trace within a preceding time window of length $\tau_2 = 3\ seconds$, i.e. considering the interval $[t - \tau_2, t]$. The relative fluorescence change was computed as
